# An ancient subcortical circuit decides when to orient to threat in humans

**DOI:** 10.1101/2023.10.24.563636

**Authors:** Hailey A Trier, Nima Khalighinejad, Sorcha Hamilton, Caroline Harbison, Luke Priestley, Mark Laubach, Jacqueline Scholl, Matthew FS Rushworth

**Affiliations:** Wellcome Centre for Integrative Neuroimaging (WIN), Department of Experimental Psychology, Tinsley Building, Mansfield Road, Oxford OX1 3TA, University of Oxford, UK; Department of Neuroscience, American University, Washington DC, USA; Department of Psychiatry, University of Oxford, Warneford Hospital, Warneford Lane, Oxford OX3 7JX, UK; Université Claude Bernard Lyon 1, CNRS, INSERM, CRNL U1028 UMR5292, PsyR2 team, Centre Hospitalier Le Vinatier, 9678 Bron, France; Wellcome Centre for Integrative Neuroimaging (WIN), Centre for Functional MRI of the Brain (FMRIB), University of Oxford, Nuffield Department of Clinical Neurosciences, Level 6, West Wing, John Radcliffe Hospital, Oxford OX3 9DU, UK

**Author notes:** **Corresponding author:** Hailey A Trier. These authors jointly supervised this work (Jacqueline Scholl, Matthew FS Rushworth).

## Abstract

Many psychiatric symptoms have been linked to threat-related perception and learning processes. In addition, however, there may also be mechanisms for balancing effectively between threat- and reward-related behaviors and these may also vary between individuals. We investigated neural activity associated with spontaneous switching between foraging for rewards and vigilance for threats with 7T fMRI. In a virtual naturalistic environment, participants freely switched between the two modes of behavior. Switching was driven by estimates of likelihood of threat and reward. Both tracking of threat and switching to vigilance were associated with specific but distributed patterns of activity spanning habenula, dorsal raphe nucleus (DRN), anterior cingulate cortex, and anterior insula cortex. Distinct distributed patterns heralded returns to reward-oriented behavior. Individual variation in DRN activity reflected individual variation in vigilance. All activity patterns were replicated in an initially held-out portion of data.

## Introduction

Anxiety and fear are prominent when mental health is poor. They are central features of psychological illnesses including generalized anxiety disorder, social anxiety, panic, and obsessive-compulsive disorder. Until recently^1^, the understanding of cognitive and neural mechanisms related to fear and anxiety has depended on experimental paradigms in which the impact of fear-inducing stimuli on behavior and neural activity can be investigated in the laboratory. Despite their elegance and continuing and critical importance^2^, such paradigms do not capture an important feature of fear-related behavior in the everyday lives of humans and other animals: the balancing of attention between threat and reward. In everyday life people must attend vigilantly to stimuli that presage adverse events but also to the stimuli predictive of positive events. To thrive they must identify and respond adaptively to stimuli predictive of positive outcomes while ensuring that their pursuit is not curtailed by dangerous threats. For the modern office worker this might mean balancing pursuit of promotion and a higher salary while avoiding difficult colleagues. For animals in natural environments the situation is analogous; daily life entails careful balancing between foraging for food while maintaining vigilance for predatory threats.

Here we focus on how people move between these two modes of behavior using a task known to reflect individual variation in core features of anxiety and depression in two large samples (discovery: N=374, replication: N=702^1^). It has been argued that considering how animals have evolved to deal with environmental threats may provide insights into fear and anxiety mechanisms both in normal and poor mental health^3–6^ and so our task adopts a similar approach here; we examine how human participants balance attention between reward- and threat-related stimuli focusing on moments of switching between the two behaviors – foraging for reward and checking for threats and the first occasion on which checking led to actual threat detection. We do this in a gamified and continuous, but carefully controlled, task in which both reward and threat stimuli were presented in a quantified manner. We measured behavior when it was predominantly guided by reward (foraging) and when it was predominantly guided by threat (checking), and in both behavioral contexts we quantified the environmental features that drove the behaviors (rate of reward and an estimate of threat proximity that we refer to as time pressure). Importantly, participants chose themselves when and how frequently to switch from one behavior to the other. In other words, our paradigm allows participants to decide when to engage in a threat- or anxiety-related response and when not to.

The type of behavior examined—freely chosen switches between threat-guided behavior and reward-guided behavior—is one important feature of the present study. The second is that we used 7T functional magnetic resonance imaging (fMRI) to identify neural processes mediating switches. The ubiquity of the need to balance reward-guided foraging and threat-guided vigilance and checking across animals suggests an evolutionarily ancient origin and its possible mediation by some of the first cephalic neural circuits evolved. These are, however, of comparatively small size in humans, often overlooked, and difficult to measure without ultra-high field imaging. We focus here on one such candidate neural circuit centered on the habenula (Hb) and dorsal raphe nucleus (DRN). While this circuit is present in primates it is unusual in that it is present in many vertebrates including even cyclostomes – jawless fish – that diverged from other vertebrates 550 million years ago^7–11^. The DRN is an important source of serotonergic innervation and like other neuromodulatory systems, such as the dopaminergic system with its origins in the ventral tegmental area (VTA) and substantia nigra pars compacta (SN), its activity is under Hb control.

While we examine activity in all four areas, an important reason for looking at DRN and Hb is that both nuclei have been linked to depression and anxiety and one way of conceptualizing such conditions is that a core feature is an inability to focus on rewarding stimuli (anhedonia) and a sustained focus on negative events. A series of recent studies have linked Hb to depression-like symptoms in rodent models^12–15^. The serotonergic system is also considered the first line pharmacological target in depression and anxiety^16–19^. As noted a previous study^1^ demonstrated the task we use here is sensitive to individual variation in anxiety; for instance, higher scores on a clinical measure of compulsive checking were reliably associated with increased vigilance and more disorganized patterns of switching between foraging and checking in the current task.

Of course, the presence of cortex in the mammalian brain suggests that Hb-DRN interactions may be influenced by cortical activity. The routes by which this might happen are, however, limited. There is little direct information about Hb connections in primates but only two cortical areas, anterior cingulate cortex (ACC) and anterior insula (AI) cortex are known to project to Hb in rodents^20–22^. We therefore also focus on these two regions of interest (ROIs). Intriguingly, in macaques, albeit in other contexts, ACC and AI carry signals related to those in DRN^23–26^. Further, AI and DRN share functional connectivity related to harm avoidance behaviors^27^ and anxiety pathology^28^. However, surprisingly little is known about these areas’ activity in many situations including predatory threat and checking behavior.

## Results

Participants were trained on the task in an online session before the scan. During the task (Fig. 1A-D) participants used arrow keys to control an animated fish in an ocean environment with rewarding food (later translated to a bonus payment), threatening predators (leading to large point loss if they ‘caught’ the fish), and a hiding space (where participants could hide from the predators). Participants attempted to gain as much reward as possible while avoiding being caught by a predator.

**Figure 1.**
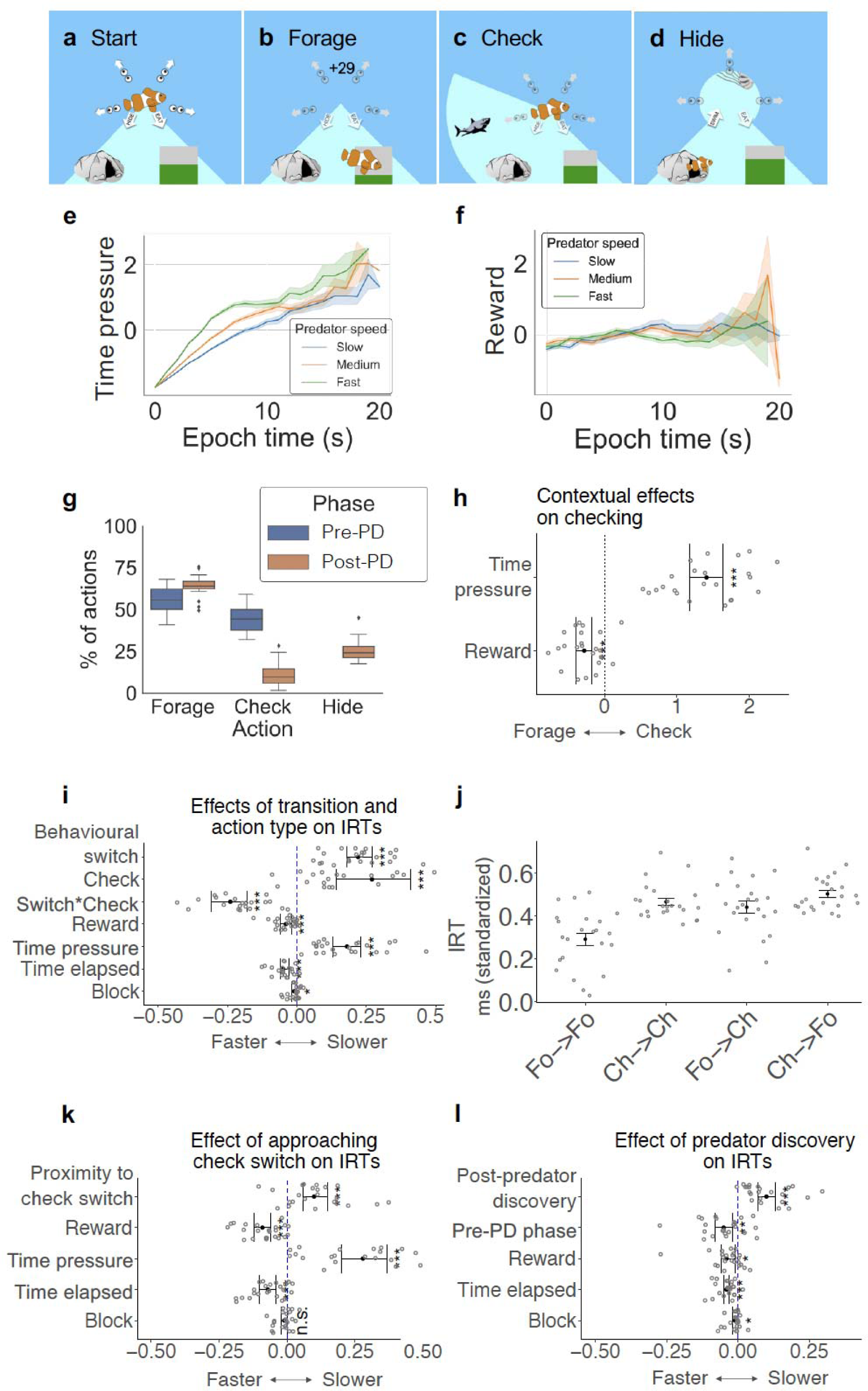
Experimental task. A) At the start of a block the fish begins in the center position. In this block the surrounding area is divided into four sections that must be searched for predators. At left (not shown here) participants see their collected energy, number of lives gained (each ‘life’=1 bar of energy), and time remaining in the current block (grey/black wheel). B) When foraging the fish dives down into the patch of food and the energy bar increases by the amount of reward gained. C) When checking, the fish checks in a particular direction. In this example the fish discovers that there is a predator in the area being checked. D) When hiding, the fish dives into a cave and is safe from the predator. It can see the predator reach the screen center and then retreat, and with another button press the fish returns to the original center position. The task lacked a traditional trial structure and participants freely chose when to forage or check. Contextual factors, however, influenced participants’ decisions, including time pressure (an index of threat level; E) and reward (F; reward appears flat because it is averaged here across all epochs); mean standardized value and 95% CI shown in 1s bins across the pre-PD phase. G-L) Contextual factors influenced pre-PD action selection (forage or check) and IRTs. G) Box plots showing action type as a percentage of all actions in each phase. H) Group mean estimates and 95% confidence intervals from two-tailed single sample t-tests showing effects of model parameters on the probability that an action was a check. I, K, L) Group mean estimates and 95% confidence intervals from two-tailed single sample t-tests showing effects of model parameters on IRTs. I) There were main effects of behavioral switch, checking, and time pressure on IRTs such that each significantly slowed reaction time (all p<0.001). IRTs were significantly faster during higher rewards (p<0.001). There was also a significant interaction between behavioral switch and checking (p<0.001). J) Group mean and standard error for the average IRTs (standardized) associated with each action sequence within each participant. IRTs were slowest when switching from checking to forage and fastest when repeating foraging. K) IRTs between forages became significantly slower as they approached an upcoming switch to checking (p<0.001). L) There was a significant main effect of whether an action occurred immediately after discovering a new predator; discovering a predator slowed IRTs (p<0.001; both pre- and post-PD phase data analyzed). *p <0.05, **p <0.01, ***p <0.001, n.s.=not significant.

In the absence of a traditional task trial structure, human participants freely and continuously chose between foraging for reward or checking for predatory threat in the virtual environment during 27 blocks each lasting 90 s. Here we focus on the two most frequent actions – foraging and checking – and the environmental variables – reward rate and an index of threat level (time pressure) that principally motivated each of the behaviors. A third action – hiding – was also available and was taken on occasions when the threat was imminent. Pressing the ‘hide’ button caused the fish to escape to a safe space where it could not be caught by the predator; a subsequent button press returned the fish to the center (Fig. 1D). Because hiding only occurred once per block, there are insufficient data for a full analysis of hiding-related behavior and neural activity.

Reward rate corresponded to the average amount of food available and followed a random walk (range: 0-90 units). Participants could always see how much food was available (proportion of green versus gray on bar on lower right of display (fig. 1A-D). When participants took the foraging action, which as we show below was more likely when reward levels were high, the fish dived down to obtain food.

Predators were hidden from participants’ view unless participants pressed a button to ‘check’ a portion of the surrounding area (Fig. 1C). At that point they were able to see a segment of the environment in which there might be a predator. Predators appeared (after a random delay, 2-10.5s) at the edge of the screen and moved toward the fish’s location at the screen center. When the predator reached the screen center it either caught the fish (causing the participant to lose one ‘life’, equivalent to 100 points) or, if the fish was in hiding, the predator simply exited the environment (ending the epoch). When participants took the checking action, if the current predator was undiscovered, the section being checked advanced clockwise to the next section of the environment at each key press; after the participant discovered the predator’s location, subsequent check responses re-checked the same location. Predator types (jellyfish, squid, shark) differed in speed, and therefore, time to reach screen center (10s, 15s or 20s). Each block had only one type of predator and participants were informed which might appear before the block start. The index of threat – time pressure – indicated the imminence of the predator’s approach (the more time elapsed after the start of the block and the faster the predator, the closer the predator would be to the center of the screen and able to catch the participant’s fish avatar (Methods, Equation 1).

Actions entailed time costs, so participants had to manage their time strategically to maximize reward. For example, each foraging action took 1.5s (see Timings in Methods). After making one foraging action, participants had to make another to obtain further reward. Alternatively, after the 1.5 s time elapsed, participants could switch to checking or hiding.

### Participants Used Task-Relevant Information to Guide Behavior

On average, participants’ actions before predator discovery (pre-PD) consisted of approximately equal numbers of checks and forages (44.68 ± 7.80% 55.32 ± 7.80% respectively; Fig. 1G) but post-predator discovery (post-PD), behavior changed; participants focused on foraging (64.01 ± 6.14%) with only occasional checks directed towards the known predator direction (10.75 ± 6.95%) and hiding actions (25.24 ± 6.15%; see Fig. 1G). Our initial analyses, therefore, examined the pre-PD phase when the two key actions, foraging and checking, were made with a similar frequency. However, we subsequently tested the post-PD data and confirmed the pre-PD results (reviewed in final figure, Fig. 8).

The moment-to-moment balance between the two behaviors – checking and foraging – was a function of the two environment features – time pressure and reward rate at the time of action (Fig. 1E, F, H). Regression analyses (Equation 4; Methods) showed that pre-PD, participants were more likely to check instead of forage as time pressure increased (*t*(22)=12.94, *p*<0.0001, *M*=1.41 ± 0.52) and as reward rate decreased (*t*(22)==-5.38, *p*<0.0001, *M*=-0.28 ± 0.25; Fig. 1H; all t-tests in Table S1).

We analyzed inter-response times (IRTs) to gain insights into when critical cognitive processes occurred. In general, IRTs were faster when time pressure increased (*t*(22)=7.30, p<0.001) and when reward level increased (*t*(22)=-4.56, *p*<0.001; Fig. 1I, Table S2). However, as noted above, as time pressure increased, participants were also more likely to check, and IRTs were slower when participants initiated checks (*t*(22)=4.23; *p*<0.001; Fig. 1I, Table S2). There was a main effect of any behavioral switch (*t*(22)=10.16), but the significant interaction between behavioral switch and checking (*t*(22)=-8.07, *p*<0.001; Fig. 1I, Table S2) demonstrated that it was switching to checking, rather than foraging, that was associated with slower IRTs. Mean IRT was slowest when switching from checking to forage and fastest when repeating forages (Fig. 1J). Such costs indicate switches between behavioral modes require cognitive resources^29–34^. (Fig. 1I; Table S3). In addition, IRTs between forages became slower as participants approached a switching to checking (*t*(22)=4.4, *p*<0.001; Fig. 1K, Table S3) suggesting participants prepared to switch to checking prior to actually making the switch. IRT was also significantly slowed by discovery of a new predator (*t*(22)=6.18, *p*<0.001; Fig. 1L, Table S4). Encountering a new threat requires additional cognitive processing relative to a check that does not reveal new information. In subsequent neural analyses we focus on understanding these behavioral switches and threat discovery points when higher IRTs indicated additional cognitive resources were deployed.

### A distributed neural network for the monitoring of threat and the transition to checking

Initial fMRI analysis focused on activity in the six ROIs (ACC, AI, Hb, SN, VTA, and DRN; Fig.2A) and examined whether it was differentially related to the two key behaviors – switching to checking and switching to foraging – and the two environmental features that drove these behaviors – time pressure and reward. Time-locking to switches in behavior allowed identification of brain activity related to these discrete events despite the free and fast nature of the task (Figs.1; S4). To examine the hypothesis that activity in these ROIs occurs during behavioral switches between foraging and vigilance, we started by analysing these ROIs alongside other brain regions by conducting a whole-brain analysis. We used GLM1 (Methods equation 6), which was structured similarly to the behavioral analyses (Methods, Equation 4), and which sought activity related to check switches, forage switches, and parametric variation in time pressure and reward rate (Fig.2). Effects of time pressure were robust enough to be apparent even in the whole brain cluster-corrected results for all ROIs (p<0.0002 two-tailed; Z>3.1; Fig. 2C, D; Table S5; Table S6). The results provide initial evidence that these brain areas might constitute a distributed circuit for orienting behavior towards potential threats. The act of switching to check was also associated with significant activation in ACC in the whole brain fMRI analysis (Fig. 2B, D; Table S5).

**Figure 2.**
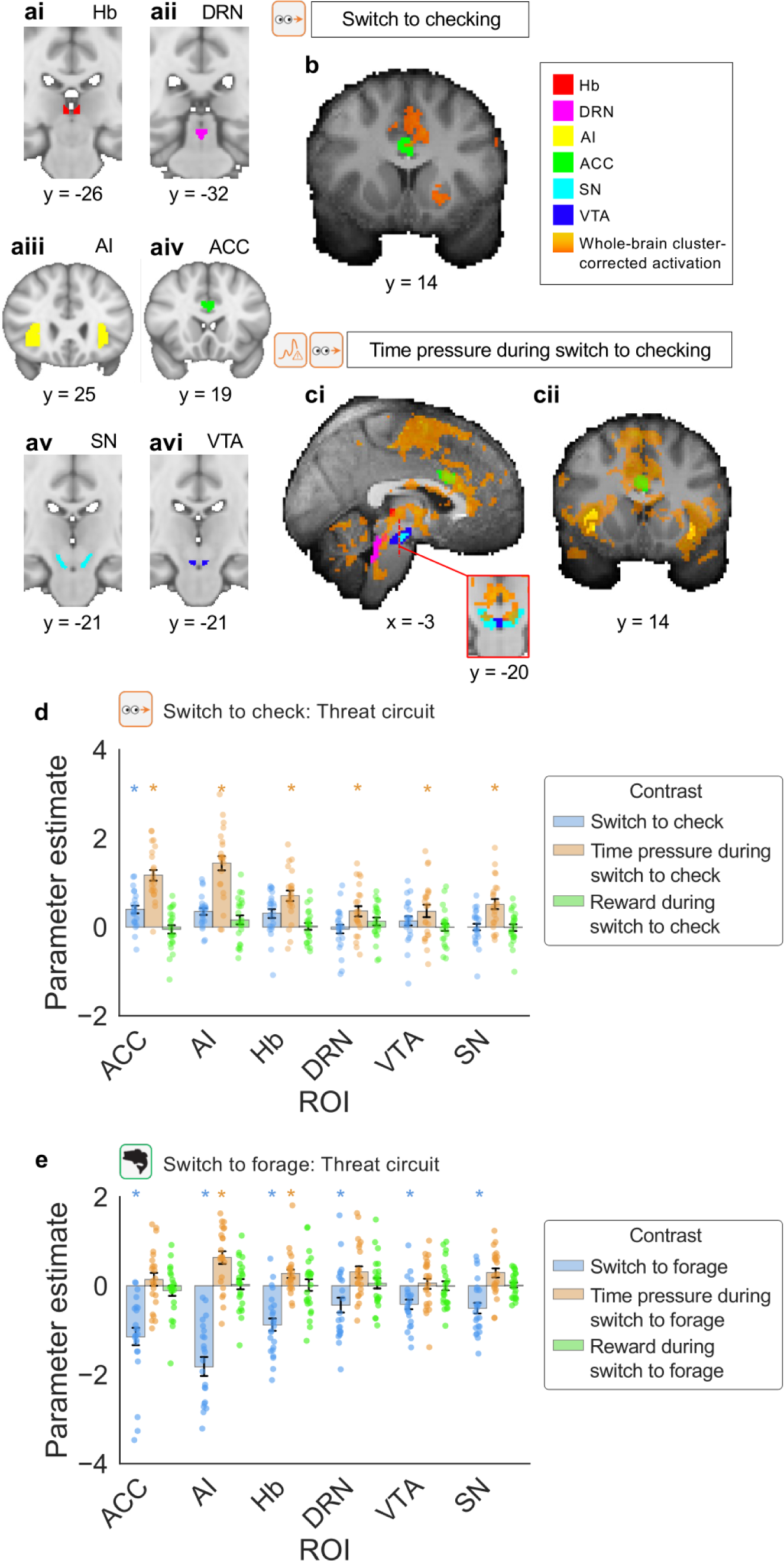
Activity related to time pressure and switches to checking. A) Activity in all six ROIs – Hb (Ai), DRN (Aii), AI (Aiii), ACC (Aiv), SN (av), and VTA (Avi) – was correlated with time pressure (an index of threat level) and, in ACC and AI activity reflected the switch to checking (p<0.0002; two-tailed; Z>3.1) . B-C) Whole-brain analysis showed that switching to check was positively associated with activity in ACC (B), and time pressure during switching to check was positively associated with activity in ACC (Ci, Cii), DRN (Ci), Hb (Ci), SN (Ci), VTA (Ci), and AI (Cii). Whole-brain cluster-based correction, Z>3.1, p<0.0001. All results are from analyses of pre-PD phase. Legend indicates color-coding of ROI masks and whole-brain cluster corrected activation. D-E) Parameter estimates (mean and standard error) in ROIs associated with threat, indexed by time pressure, reward level, and effects of check or forage switches. D) Analyses of ROI activity reveals activity in all cases encodes check switches and time pressure. E) The same areas are not activated by reward level and they are deactivated by forage switches. n *=p < 0.0001 after whole-brain cluster correction.

To confirm our interpretation of the fMRI data we extracted parameter estimates (*β* weights from GLM1) in the ROIs shown in Fig. 2A. We examined parameter estimates linked to the first instance of each behavior – foraging or checking – on each occasion that participants switched (referred to, respectively, as forage switch and check switch, modeled as constants. In addition GLM1 identified parametric variation in the two key environmental features that motivated the types of switching, time pressure and reward level during each of the two switch types. A three way ANOVA performed on the parameter estimates revealed that across the six ROIs there was a main effect of switch type (switching to checking was associated with more effects than switching to foraging: F(1, 528)=30.03, *p*<0.0001; blue bars are more positive in Fig.2d versus 2e); a main effect of type of environmental feature (time pressure effects were stronger than reward rate effects: F(1, 528)=114.72, *p*<0.0001; ochre bars are higher than green bars in Figs.2d and 2e) and a three-way interaction between ROI, switch type, and environmental feature suggested time pressure signals were stronger than reward rate signals especially when switching to checking and this was particularly true in some brain areas (ACC, AI, Hb: F(5, 528)=2.33, *p*<0.0001; ochre bars are especially larger than green bars on the left in Fig. 2d versus 2e). In summary, the ANOVA revealed that time pressure-related activity, as opposed to reward rate modulations, were most apparent when switching to checking as opposed to foraging and this was especially true in ACC, AI, and Hb (Fig. 2B-E). Moreover, it was apparent that activity related to reward rate was negligible even at the time of forage switches in these areas (apparent in the near zero reward effects in Fig. 2D and in the absence of any significant effect of reward at the time of checking in the whole brain analysis; Fig. 2A). Below we discuss reward rate-related activity at the time of switching to foraging that was found in other structures, but at the time of checking the absence of significant, univariate reward-related activity in the ROIs suggested a strong attentional focus on time pressure.

So far the analysis had suggested that all six ROIs carried time pressure-related information (apparent in the significant effects of time pressure in the whole brain analysis; p<0.0002; two-tailed; Z>3.1 mentioned above). It is well established that some of these areas, such as the cortical regions ACC and AI carry a range of additional signals in a diverse range of tasks. We were, however, interested in the possibility that one of the subcortical areas such as DRN, where activity-behavior correlates are less well established, might be especially concerned with the tracking of this variable in order to bring about a change in behavior. This might be apparent if the strength of the time-pressure signal were especially closely related to the likelihood that a switch might occur. To examine this possibility we tested whether individual variation in time pressure signal across participants was predictive of individual variation in the frequency of checking as indexed by the percentage of responses that were checks. Not only did we carry out this test in DRN but, for comparison, in the three other subcortical areas. There was indeed a relationship between the strength of the time pressure signal and checking frequency across participants (Pearson’s r = 0.52, *p* = 0.012 Fig. 3). This remained true even after Bonferroni correction for multiple tests conducted across the four subcortical areas. The relationship between the time pressure signal strength at the time of switching to checking is not a mandatory one that is found in all brain areas in which there is a time pressure effect; while reliable effects were not found in other areas (all *p* > 0.05), the relationship in the DRN was replicated in an initially held portion of the data (from the period in each block after the predator was discovered; this and other replication tests are discussed below and in Figs. 3B and 7). Notably, individual variation in the balance between foraging and checking has been shown to be related to individual variation in clinical indices of compulsive behaviors, such as checking, in a large sample study^1^. The present results suggest that individual variation in DRN activity might be related to checking including when it becomes compulsive and problematic.

**Figure 3.**
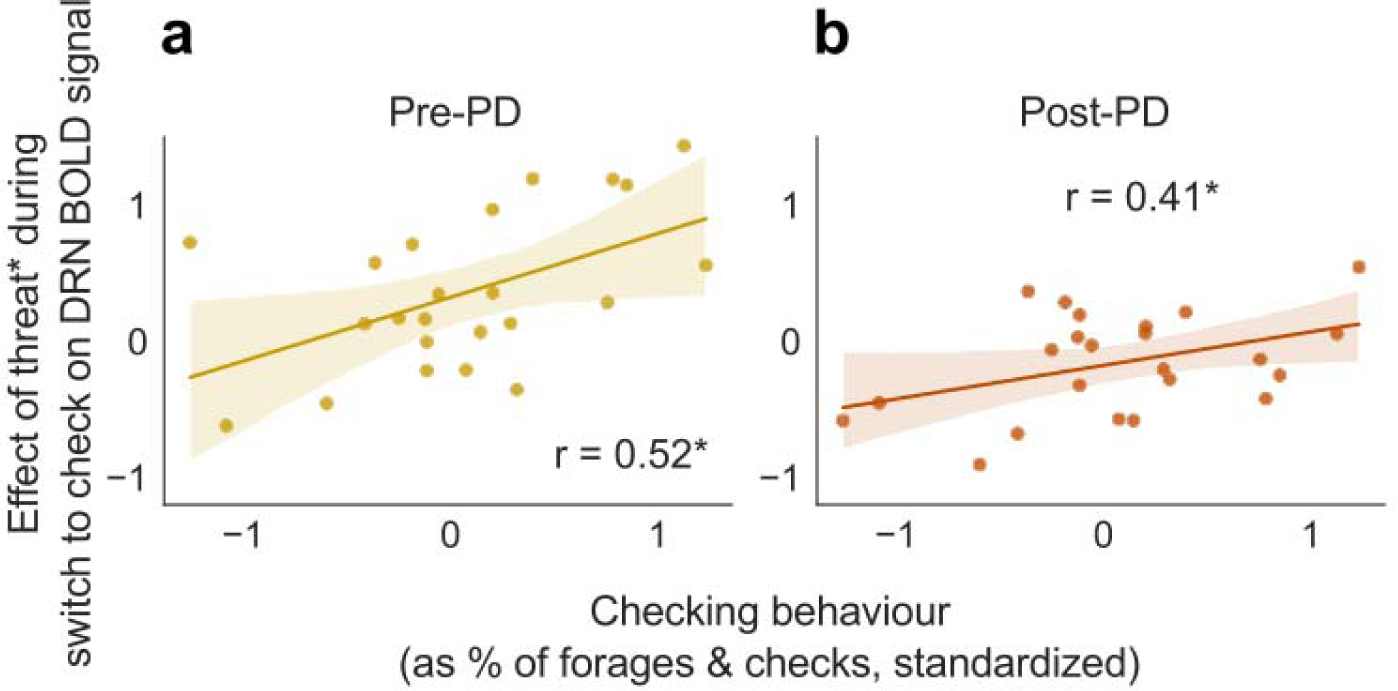
BOLD signal in DRN corresponding to threat level (in pre-PD, indexed as time pressure; in post-PD, indexed as proximity) during switching to check was significantly related to checking behavior as measured by checks as a percentage of forages and checks combined. The pre-PD correlation (Pearson’s r=0.52, p = 0.012) replicated in post-PD behavior (Pearson’s r=0.41, p = 0.049). *=p<0.05.

While we focus here on ACC, AI, Hb, SN, VTA, and DRN, it was also apparent from the whole brain analysis (Fig. 2; Table S5) that time pressure during checking and the act of switching to checking was associated with activity in superior colliculus, pulvinar nucleus of the thalamus, and dorsal and ventrolateral parts of the periaqueductal grey. This is consistent with the suggestion that these brain structures may mediate fast responses to threat stimuli^35^.

### Interactions across the distributed network for threat monitoring and transition to checking

Next, we sought to understand how the DRN interacted with the other areas to encode time pressure and to bring about the process of behavioral change when participants switched to checking. We therefore used psychophysiological interaction (PPI) analyses^36^ (Methods, Equation 8) to examine relationships between DRN (in which the link with behavior is prominent; Fig.3) and three other areas, ACC, AI, and Hb, because, as noted above: i) their time pressure signals were stronger than their reward rate signals especially when checking and their switching signals were especially different when checking as opposed to foraging (Fig. 2D, E); ii) their anatomical connections suggest they are especially well placed to influence DRN^7–11,20–22;^ iii) in macaques, ACC and AI carry signals related to those in DRN^23–26^. We found two types of interactions. The first occurred as a function of switching from foraging to checking. The second also occurred as a function of switching from foraging to checking but, in addition, this interaction also varied with time pressure.

The first pattern of interaction was evident between ACC and Hb (two-sided Wilcoxon signed rank test, Z=3.28, *p*=0.001; Fig. 4Aii); full PPI analysis results reported in Table S7). Stronger ACC activity was associated with stronger Hb activity during check as opposed to forage switches (Figs. 4Aiii and 4Aiv illustrate how ACC-Hb interactions differed depending on switch direction). Notably, the average peak time of this effect was shortly prior to the first checking button press. Given that fMRI peak effects are delayed by the hemodynamic response function, it reflects neural events occurring several seconds before the actual check switch. This is consistent with behavioral evidence (Fig. 1K) showing participants prepared to switch to checking several seconds before actually switching. This pattern is specific to these areas; no evidence was found for similar interactions involving AI and DRN (Fig. S1A-C). To demonstrate the specificity of the ACC-Hb pathway we carried out a factorial style analysis examining interactions between two cortical regions, ACC and AI, and two subcortical regions, Hb and DRN. A two-way ANOVA revealed a main effect of cortical region (ACC and AI) on the extent to which switching to check moderated functional connectivity with subcortical ROIs Hb and DRN (F(1, 180)=7.03, *p*=0.009); Fig. 4Ai; Table S3); post-hoc Tukey HSD test found that peak values were greater on average for ACC than AI (*p* adj.=0.012; 95% CI=[-0.04, -0.01]; Table S8). There was also a significant interaction between cortical and subcortical region (two-way ANOVA, F(1, 180)=16.13, *p*<0.001; Fig. 4Ai; Table S9), with mean peak values being highest for interaction between ACC and Hb as a function of checking which, as reported above, was significant when tested in isolation.

**Figure 4.**
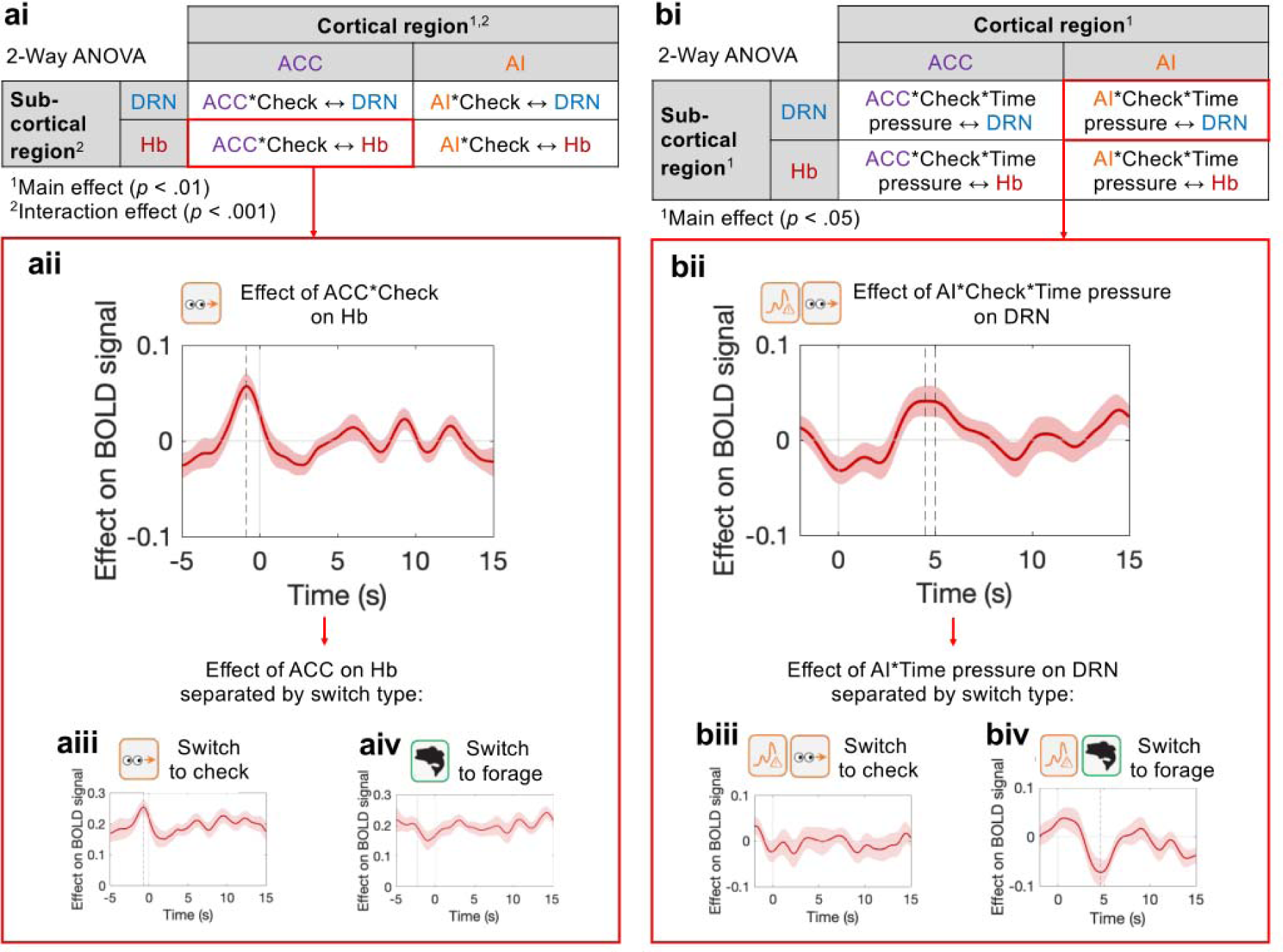
Interactions between cortical areas ACC and AI and subcortical areas Hb and DRN were modulated as a function of switching to checking (A) and as a function of both switching to checking and time pressure (B). Ai) Two-way ANOVA showed a main effect of cortical region (ACC/AI; p<0.01) and an interaction between cortical and sub-cortical region (Hb/DRN; p<0.001) on the extent to which check switches moderated functional connectivity. Average peak values were highest for ACC and Hb activity moderated by checking. Other interactions tested in the ANOVA are plotted in Figure S1A-C. Aii) Functional connectivity between ACC and Hb was significantly moderated by switching to check (p<0.01). ACC activity interacted with Hb activity differently during check switches (Aiii) versus forage switches (Aiv). Bi) Two-way ANOVA showed main effects of both cortical (p<0.05) and sub-cortical (p<0.01) ROIs on the extent to which check switch and time pressure moderated functional connectivity. Average peak values were highest for interactions with AI as the cortical ROI, and for interactions with DRN as the sub-cortical ROI. Other interactions tested in the ANOVA are plotted in Figure S1D-F. Bii) A three-way interaction between AI activity, check switch, and time pressure moderated DRN activity (p<0.05). AI and time pressure affected DRN activity differently during check switches (Biii) versus forage switches (Biv). All analyses shown used pre-PD phase data. Significance testing on time course data was performed using a leave-one-out procedure on the group peak signal. Dashed line indicates the average time of peaks across which two-sided Wilcoxon signed rank test was significant. Absence of dashed line indicates non-significance.

The second pattern of interaction, encoding of time pressure and switching to check between AI and DRN, is summarized in (Fig. 4Bii). Like the first interaction, it occurred between a cortical and subcortical region as a function of switching to checking versus switching for foraging but now the interaction was between AI and DRN and it occurred as a function of an additional factor – *time pressure* – the environmental variable that drove participants to switching. There was a significant three-way interaction between AI activity, time pressure, and switching to check that modulated DRN activity (Z=1.98, *p*=0.048; Fig. 4Bii, Table S7), such that stronger AI activity was associated with stronger DRN activity as a function of switch to checking and as a function of *time pressure* (see Fig.3Biii, 3Biv for depictions of how AI activity and *time pressure* were related to DRN activity differently depending on the direction of switch). The interaction reflects a relative decrease in AI-DRN coupling as a function of time pressure when participants switch to foraging. Again, this pattern of activity exhibited a degree of specificity (Fig. S1D-F); a factorial analysis in which we examined interactions between the two cortical regions, ACC and AI, and the two subcortical regions, Hb and DRN. There was a main effect of sub-cortical ROI (Hb and DRN; two-way ANOVA, F(1, 180)=11.11, *p*=0.001; Table S10) and a main effect of cortical ROI (ACC and AI; F(1, 180)=4.22, *p*=0.04; Table S10) on the extent to which switching to check and time pressure moderated functional connectivity between the four regions. Although, the interaction term failed to reach significance, post-hoc Tukey HSD tests found that peak values were on average greater for analyses with DRN as the sub-cortical ROI (*p* adj.=0.001, 95% CI=[-0.06, -0.01]; Table S11) and greater for analyses with AI as the cortical ROI (*p* adj.=0.047, 95% CI=[0.0003, 0.05]; Table S12).

### Habenula interactions with dopaminergic and serotonergic nuclei during re-orienting to threat stimuli

In the previous section we identified a route of interaction between cortex (AI) and DRN and a route of interaction between cortex (ACC) and Hb. The Hb is, however, itself an important source of influence over DRN, SN, and VTA^37^. In the next analyses we examined how Hb interacted with DRN and compared the pattern we found with SN and VTA at the time that checking or foraging actions were initiated. PPI analyses revealed Hb exerted a specific effect, in the sense that it was related to time pressure, but a general effect in the sense that the impact in DRN was similar to those seen in VTA and SN; in each case, greater Hb activity was associated with greater activity in the target brain region as a function of both checking and time pressure. This effect was specific to check switches but found in the interaction patterns of Hb with all three areas: two-way ANOVA showed a main effect of switch type (check versus forage) on the extent to which the effect of Hb activity was moderated by time pressure in all three brain areas (F(1, 270)=4.94, *p* = 0.027; Table S13, Fig. 5A), with higher peak values associated with checks (Table S14). The average time at which peaks were significant for each PPI analysis for the three areas (DRN, VTA, SN) was 5.79s ± 0.54s after the time-locking event – the initiation of the checking action, suggesting (once the BOLD haemodynamic response is taken into account) that this effect occurred after the ACC-Hb interactions illustrated in Fig.3 and was elicited at the onset of the checking action itself. Significance testing of peak values in separate PPI analyses confirmed functional connectivity with Hb was moderated by switching to check, time pressure, and activity in SN (Z=2.16, *p=*0.031; Fig. 5B), VTA (Z=2.13, *p=*0.033; Fig. 5C), and DRN (Z=2.04, *p=*0.042; Fig. 5D).

**Figure 5.**
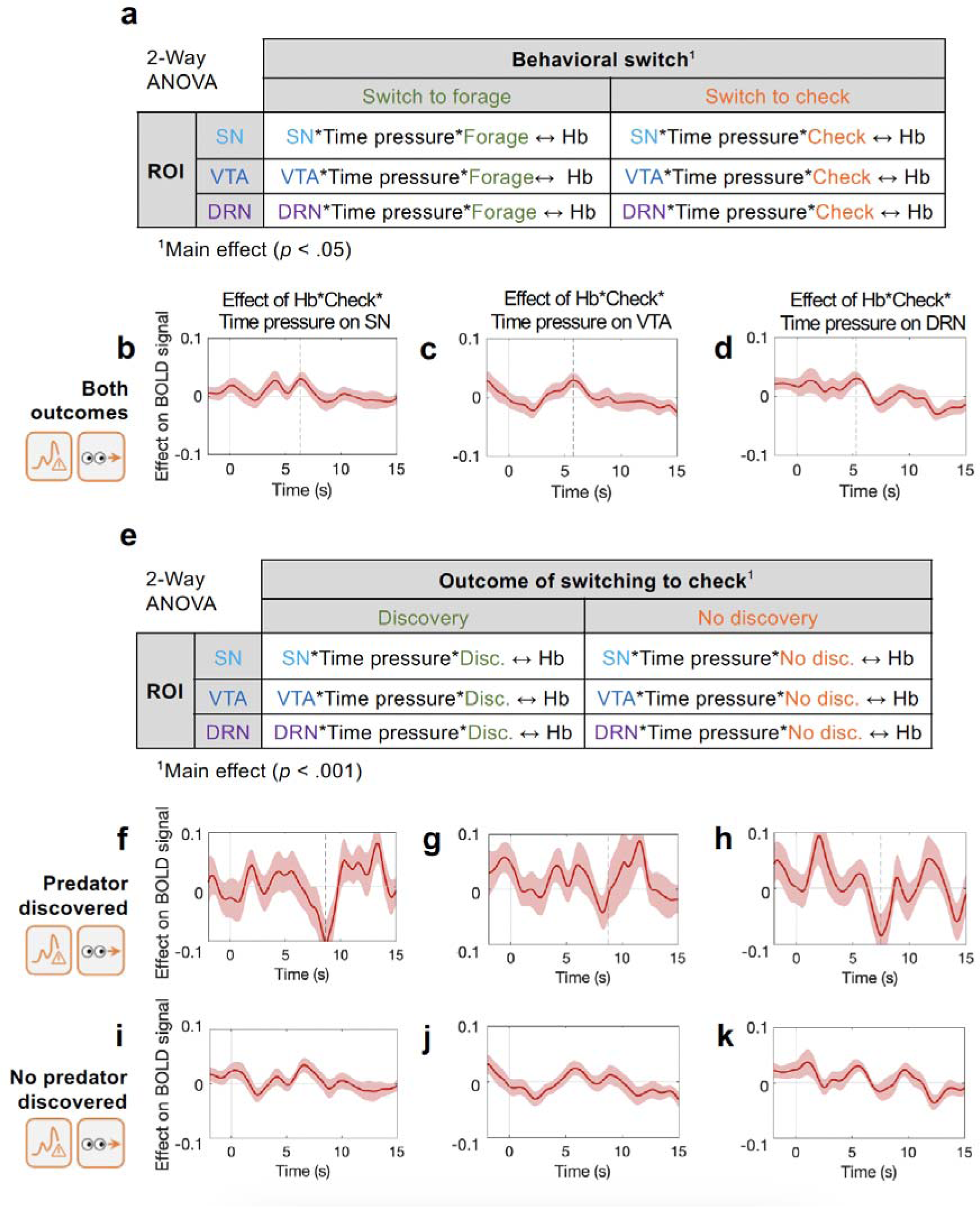
Functional connectivity between Hb and SN, VTA, and DRN were affected by time pressure, switching to check, and check outcome (predator discovery/non-discovery). A) Two-way ANOVA showed a main effect of check outcome (p<0.001) on the extent to which switching to check and time pressure moderated functional connectivity with Hb. B-J) PPI analyses showing interaction between Hb and SN (B, F, I), VTA (C,G,JI), and DRN (D,H,KJ) as a function of time pressure and checking. When we looked at all check trials (B, C, D) there was a three-way interaction between Hb activity, time pressure, and switching to check on B) SN, C) VTA, and D) DRN activity; as time pressure increased, and when participants switched to checking, then higher Hb activity was associated with higher activity in SN, VTA, and DRN. Subsequent analyses looked at checks separately as a function of whether they revealed a predator for the first time (F, G, H) or not (I, J, K). When a switch to checking revealed a new predator, there was an additional, slightly later, negative three-way interaction between Hb activity, time pressure, and switching to check in SN, VTA, and DRN. The timing of positive interaction highlighted in panels B-D links it to the checking action itself while the later timing of the negative interaction in panels F-K links it to predator discovery. Significance testing on time course data was performed using a leave-one-out procedure on the group peak signal. Dashed line indicates the average time of peaks across which two-sided Wilcoxon signed rank test was significant.

Several features of the results are notable. First, that Hb interacted with ACC as a function of check switches and with VTA, SN, and DRN as a function of both time pressure and check switches suggests that Hb may play a unique role in facilitating communication between cortical and sub-cortical areas to guide behavioral change. Indeed, further PPI analysis showed that activity in Hb was moderated by an interaction between AI activity and time pressure, regardless of switch type (Z=2.16, *p*=0.031; Table S7; Fig. S5). Second, Hb interactions with DRN, VTA, and SN were all similar. By contrast, the AI-DRN interaction during check switches reported above (Fig. 4Bii) was specific to those two areas.

### Subcortical regions encode threat discovery

The previous section considered proactive behaviors to potential threats as participants chose voluntarily when to initiate checking for predators regardless of whether or not the checks led to predator discovery. Voluntary switches in behavior are a key aspect of the current task but previous studies of Hb and interconnected structures have focused on very different aspects of its activity when surprising events are encountered^37^. Therefore in the next section, we focus on the reactive aspect of threat monitoring when participants made a checking response that actually led to the new discovery of the predator. We did this by continuing to employ the same index of threat – time pressure – but while previously we had considered both checks that led to predator discovery for the first time in a block and checks that did not, we now compared these two types of checks (we refer to this as the *check outcome* factor). As noted already, many checks did not lead to predator discovery but discovery checks entailed surprise and slowing of IRTs (Fig. 1K). Because Hb and some of the regions that it projects to (the dopaminergic midbrain) have been reported to carry signals relating to surprising events, such as reward prediction errors^11,38^ and because there are similar projections from Hb not just to dopaminergic midbrain but also DRN^37^, we examined the interactions between Hb and SN, VTA, and DRN using a PPI approach as above. Now, however, in addition, the interactions were examined as a function of whether checking led to predator discovery. Interactions were indeed sensitive to check outcome. Two-way ANOVA showed a main effect of predator discovery (F(1, 132)=23.77, *p*<0.0001; Fig. 5E, Table S15) on peak values for the three-way interaction between Hb activity, switching to check, and *time pressure* with all three areas (SN, VTA, and DRN). Notably, the average time at which peaks were significant for each PPI analysis was 8.25s ± 0.71s after button press (Fig.5.f-h); given hemodynamic lag, this suggests that the three-way interaction between switching to check, time pressure, and predator discovery/non-discovery on interactions between Hb and the others areas was elicited by an event occurring at the point of predator discovery/non-discovery rather than at the initiation of checking (compare Fig.5f-h with Fig.5b-d).

Further analyses focusing either on predator discovery events or predator non-discovery events revealed predator discovery was linked with relatively inhibitory relationships between Hb and each ROI (SN, VTA, and DRN) as a function of time pressure. For these tests, as before, we examined activity time-locked to the initiation of checking but instead of looking at the relatively early activity actually linked to the initiation of the check (Fig.5B-D), we focused on later activity, approximately 9s after check initiation, when the participants had seen the outcome of checking. In this way we were able to look at how activity differed when the check outcome was predator detection (Fig.5F-H) or absence (Fig.5I-K) PPI results showed that when switching to checking resulted in predator discovery, functional connectivity with Hb was moderated by a three-way interaction with switching to check, time pressure, and SN activity (Z=-2.43, *p*=0.015; Fig. 5F), VTA (Z=-2.95, *p=*0.003; Fig. 5G), and DRN (Z=-2.10, *p=*0.036; Fig. 5H). In each case greater Hb activity was associated with lower activity in the region of interest. This is consistent with evidence that Hb has an inhibitory effect on midbrain dopamine neurons in the VTA and SN^15,39,40^ in response to negative-valence and aversive stimuli^12,15^, and also plays a role in regulating DRN serotonergic neurons^41^. No such relationship was found for any target region when switching to check did not reveal a threat (Fig. 5I-K).

### A distributed neural network for the monitoring of reward and the transition to foraging

So far, we have focused on proactive switching from foraging to checking as a function of the potential for threat – *time pressure* – and as a function of reactive detection of threat when the predator was discovered. We next looked for evidence of complementary neural activity mediating behavioral switch in the opposite direction – from checking to foraging. When looking at forage switches, we also considered reward rate because switching from checking to foraging was promoted by higher reward rates (Fig. 1H). The behavioral analyses of IRTs (Fig. 2) had suggested that foraging was the participants’ default behavior during the task but that this behavior was intermittently interrupted by checking.

As already noted, the network of areas linked to time pressure and check switches exhibited little reward rate-related activity during checking and was less active during switches to foraging as opposed to checking (Fig. 2B, D, E). This is striking given that the actions participants made to forage or check were nearly identical finger movements. The difference in the goal of the action—foraging or checking— despite similarity in the nature of the finger movement meant that the distributed pattern of activity across the brain was profoundly different.

However, it was possible to find reward rate-related activity when participants were switching to foraging. This was apparent in significant parametric effects of reward rate at the time of switching to foraging in the whole brain analysis (GLM1; Fig. 6; Table S5). These were prominent in a relatively ventral part of the left striatum either side of the internal capsule and in the cross bridges spanning it (Fig. 2bi). Additionally, increases in activity related to forage switches and reward levels were found in adjacent parts of the precentral gyrus in dorsal premotor and motor cortex. The reward-related and forage switch related activity is also summarized in ROIs at each location (Fig. 6C, D). In a final analysis we looked at interactions between these areas as a function of reward rate and the forage switches as opposed to check switches. There was a positive interaction between precentral gyrus and striatum as a function of reward rate that increased further when this led to a switch to foraging (Z=1.98, *p=*0.048; Fig. S3A); this occurred at the time of button press and was thus likely elicited by preparation for the switch to foraging.

**Figure 6.**
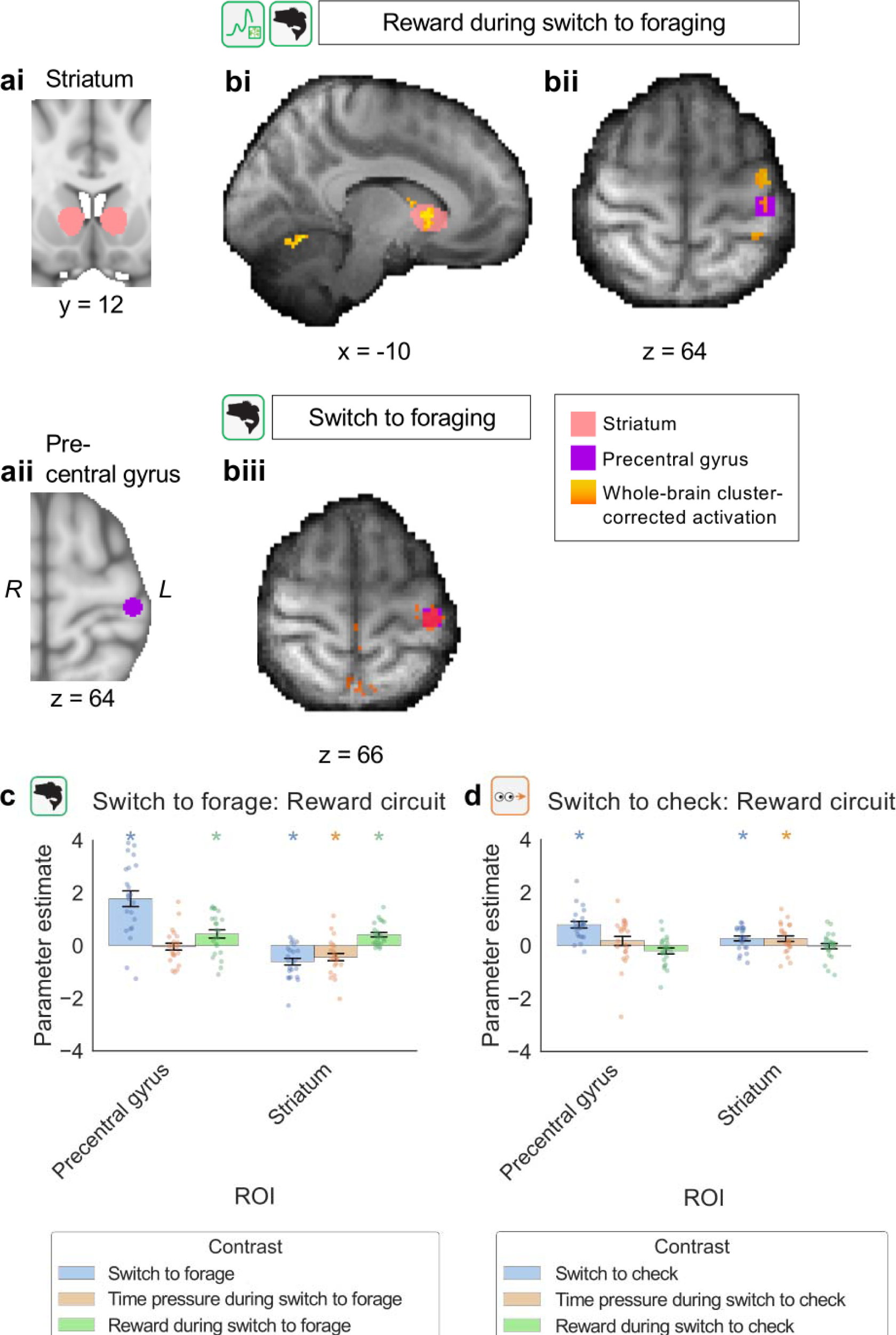
Activity related to switching to foraging, time pressure, and reward. A-B) Whole-brain analysis revealed activity in the striatum (Ai, Bi) and precentral gyrus (Aii, Bii) was correlated with reward during switching to forage (B). In addition, activity in the precentral gyrus was positively correlated with the act of switching to forage itself (Biii). Whole-brain cluster-based correction, Z>3.1, p<0.0001. Legend indicates color-coding of whole-brain cluster corrected activations and ROI masks. C-D) Parameter estimates (mean and standard error) associated with switches to foraging and checking as well as environmental variables (time pressure, reward) during each switch. There was a positive reward signal in the precentral gyrus and striatum during switches to foraging (C) that was greater than the equivalent in switches to checking (D). Significance testing on time course data was performed using a leave-one-out procedure on the group peak signal. Dashed line indicates the average time of peaks across which two-sided Wilcoxon signed rank test was significant. *= p < 0.0001 after whole-brain cluster correction.

**Figure 7.**
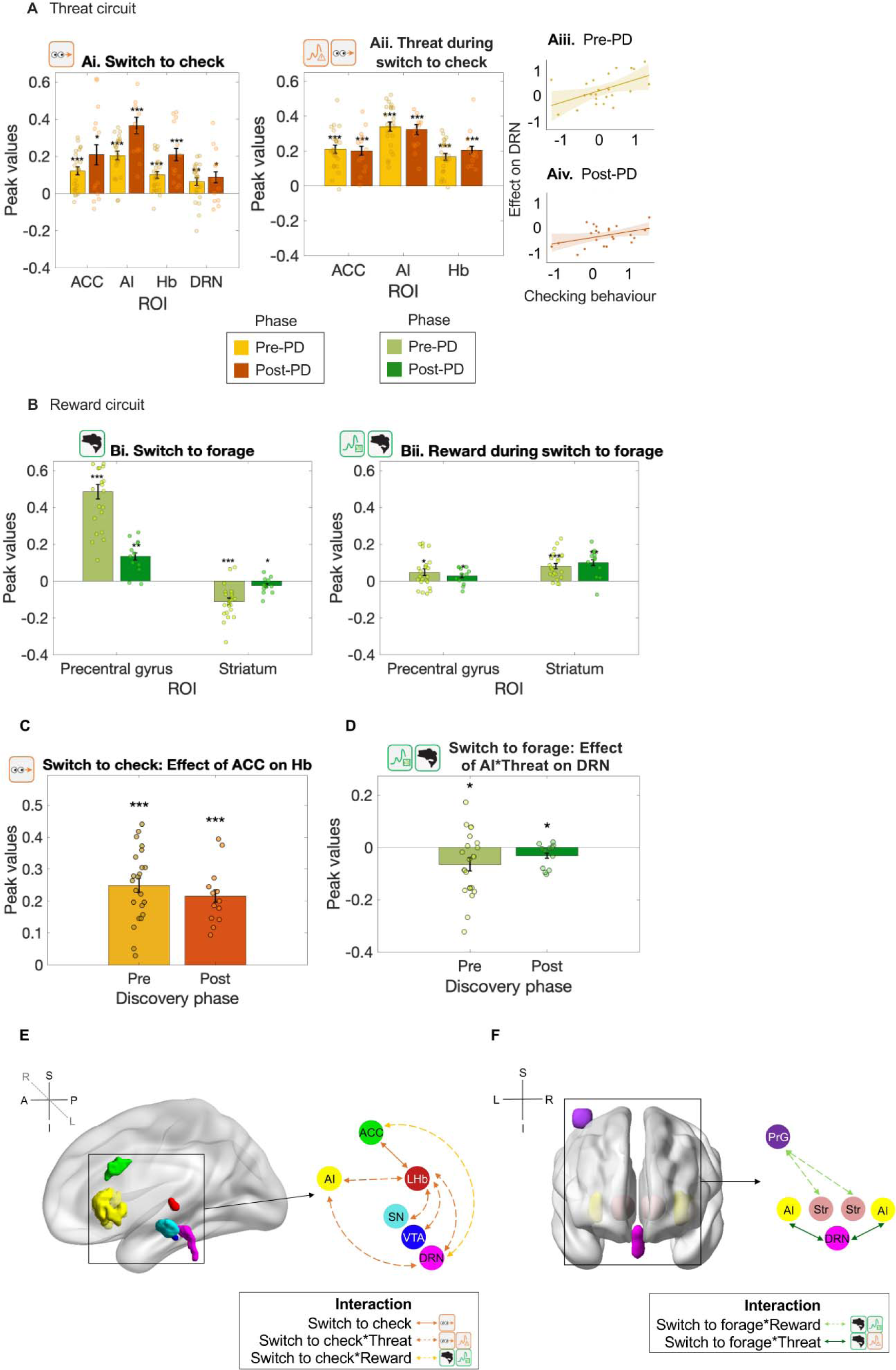
Replications in post-PD phase. Pre-PD phase effects replicated in post-PD phase data. A) As in pre-PD phase results shown in Fig. 2C, post-PD BOLD activity in Ai) ACC, AI, Hb, and DRN encoded switching to check, and Aii) ACC, AI, and Hb encoded threat level (in pre-PD, indexed as time pressure; in post-PD, indexed as proximity) during switching to check (all p < 0.05). As shown in Fig. 2, BOLD signal in DRN related to time pressure during switching to check was correlated with checking behavior (as % of all checks and forages) in both pre-(Aiii) and post-PD (Aiv) phases (both p < 0.05). As in pre-PD phase results shown in Fig. 6C, post-PD BOLD activity in Bi) precentral gyrus and striatum encoded switching to forage, and Bii) reward level during switching to forage (replication in post-PD period tested in the same voxels identified as significant in the prediscovery phase; all p < 0.05). Key PPI results also replicated: C) as in Fig. 4Aiii, functional connectivity between ACC and Hb was mediated by switching to check; D) as in Fig. 4Biv, an interaction between time pressure and AI activity moderated DRN activity; as in the pre-PD phase this was due to a negative interaction between AI activity and time pressure on DRN activity during switching to forage (all p < 0.05). In summary, a distributed neural network encodes E) time pressure and switching to check, and F) reward level, time pressure, and switching to forage.

#### Replication in post-PD phase

To replicate our findings, we performed the same analyses in the post-PD phase. Replications of the results shown in figures 2-4 and 6 are shown in a condensed form in Fig 7). Because no time analogous to the point of predator discovery existed in the post-PD phase, it was not possible to examine activity patterns such as those shown in figure 5. Individual variation in the DRN threat signal (pre-PD: indexed by time pressure, post-PD: indexed by proximity) continued to be correlated with individual variation in checking (Figs. 7a Iii-Aiv) although the main effect of the same signal was not identifiable when no consideration was taken of check rates. This result underlines the close link between DRN threat signals and checking behavior. All other effects, including PPI results, were replicated (Fig. 7; Table S16).

## Discussion

Life in natural environments requires humans and other animals to strike a balance between reward pursuit and threat monitoring. While progress has been made in understanding mechanisms underlying reward- and threat-guided behavior independently, how people and other animals spontaneously switch between the two behavioral modes is not well understood. In addition to the fundamental importance of such decisions, individual variation in how they are taken is related to individual variation in clinical anxiety scores such as compulsivity^1^. In the present experimental paradigm, participants freely managed their time and decided themselves when to switch between foraging for rewards versus checking for approaching predators. Our analysis focused on switching points where participants spontaneously changed between foraging and checking. We initially focused on the task phase before participants discovered the predator (pre-PD), because during this phase switches to checking and foraging occurred with approximately equal frequency. However, effects found in the pre-PD phase were subsequently replicated in the post-PD phase (Fig. 7).

We found evidence of activity and specific interaction patterns across a distributed circuit (Figs. 2-7). DRN activity tracked time pressure, a measure of the imminence of the threat and the strength of the signal was closely related to whether a switch to checking occurred (Fig. 3; 7). Robust patterns of interaction were also found between DRN and Hb and ACC as a function of either switching to checking or time pressure (which promoted checking; Figs4,5). AI-DRN interactions were especially specific and occurred as a function of both time pressure and witching to check. Once the hemodynamic response function is taken into account, the early timing of interaction effects—peaking less than 5s after switching to checking —indicates that the neural signals preceded the switches to the vigilant state.

These specific interaction patterns were complemented by a second set of more general interaction patterns, observed between Hb, on the one hand, and, on the other hand, not just with DRN but also with SN, and VTA. Functional connectivity between Hb and DRN/VTA/SN was mediated by threat level and check switches in a way that was sensitive to check outcome (i.e., whether the check revealed a new threat or not). Initially when checks were made, interactions were positive. However, once a new threat was discovered, a strong negative relationship between Hb and each region was observed as a function of threat level and checking (Fig. 5). The inhibitory influence exerted by Hb over VTA/SN dopaminergic and DRN serotonergic neurons has previously been emphasised^12,15,39–41^. There may initially be a small but significant risk-related excitation between Hb and DRN, VTA, and SN as a function of time pressure when the check is made; some VTA/SN neurons exhibit excitation that varies with risk during the time between a reward-predicting stimulus and an outcome^42–44^. DRN activity has also been reported to reflect reward variance^45^. The excitatory neurotransmitter glutamate has been reported in Hb afferents to DRN^41^.

One influential idea has been that DRN encodes a prediction error signal in response to aversive stimuli that might complement one for tracking rewards in dopaminergic nuclei^46–48^. More recent research, however, has suggested that while DRN activity may indeed reflect aversive events it also reflects appetitive ones^24,45,49–52^ and that a prolonged increase in serotonin increases learning signals for aversive and appetitive stimuli^53^. Nevertheless, the induction of activity in DRN does not appear to be rewarding in a simple way but instead, it leads to changes in the choices animals make as a function of both costs and benefits; for example, after DRN stimulation, mice become more likely to wait through a delay to obtain a reward^50,51,54–61^. The current results suggest that this is because DRN is not simply concerned with regulation of patience or motor inhibition when deciding how long to wait for a reward but, instead, that it has a more general role in regulating the impact that aversive and appetitive prospects have on behavior. One possibility, therefore, is that DRN has a general role in tracking both rewarding and aversive features of the environment, with the latter especially salient in the current study, in order to redirect the motivational focus for behavior. In the present study, redirection occurs between reward- and threat-related motivations. In another recent, unpublished study from our laboratory we have observed DRN activity as macaques switch between reward-related motivation and inaction. Relatedly activity patterns in zebrafish DRN can be interpreted as being related to switches between reward-guided motivation and exploration^62^. The current results suggest these insights from fish and non-human primate studies may be useful in understanding how humans decide when and how frequently to direct behavior to threat as opposed to rewards. In addition they shed light on how cortical and sub-cortical regions work together to track behaviorally relevant stimuli and to switch motivational modes between threat vigilance and other reward-related motivations not just in healthy behavior but potentially also when vigilant behaviors, such as compulsive checking, become overwhelming in frequency and potentially clinically problematic.

## Methods

### Subjects

24 healthy adult participants (15 females), aged 18 to 35, completed the study. Participants were paid £10 and £15 per hour for the online and scan sessions respectively, plus a performance-dependent bonus (£5.10 ± 0.86). Ethical approval was given by the Oxford University Central University Research Ethics Committee (Ref-Number MSD-IDREC-R55856/RE006). One participant was excluded from all analyses because they did not make enough ‘check’ actions to compute all regressors of interest in the model. Behavioral data from all other participants were included in analyses.

### Task

We designed a gamified foraging task in which participants freely made a series of choices with the goal of earning as much money as possible. During the task (Fig. 1 A-D) participants used arrow keys to control an animated fish in an ocean environment where there was rewarding food (later translated to a bonus payment), threatening predators (leading to large point loss if they ‘caught’ the fish), and a hiding space (where participants could hide from the predators). Participants chose freely between three actions: ‘forage’ for food, ‘check’ for predators, and ‘hide’ in a safe space. Participants were trained on the task in an online session before the scan. Each participant received one of two task schedules, each having the same scheduled blocks but in a different, randomised order.

#### Forage

The average amount of food available varied randomly per second (range: 0-90 units). Participants could always see how much food was available. When foraging the fish dived down to obtain food (translated later into money; see Fig. 1B).

#### Check

Predators were hidden from participants’ view unless participants pressed a button to ‘check’ a specific portion of the surrounding area (Fig. 1C). Predators appeared (after a random delay, 2-10.5s) at the edge of the screen and moved toward the fish’s location at the screen center. When the predator reached the centre of the screen it either caught the fish (causing the participant to lose one ‘life’) or, if the fish was in hiding (see below), the predator quickly exited the environment (ending one ‘predator epoch’). If the current predator was undiscovered, the section being checked advanced clockwise to the next section of the environment at each key press; after the participant discovered the predator’s location, successive key presses re-checked the same location. Predator types (e.g. shark) differed in speed (10s, 15s or 20s to reach screen center) (Fig. 1E, F). Each block (see below) had only one type of predator, appearing one at a time, and participants were informed which before the block start.

#### Hide

Pressing the ‘hide’ button caused the fish to escape to a safe space where it could not be caught by the predator; a subsequent button press would return the fish to the center (Fig. 1D).

#### Blocks

Each of 27 experimental blocks lasted 90s and had a different combination of (1) predator type (i.e. predator speed, three levels, see above) and (2) number of segments in which participants could check for predators (range 1-4). With fewer segments, more of the environment was visible during each check action and therefore fewer checks were required to survey the entire surrounding area. Each participant received one of three schedules, each having block and reward conditions randomly generated with the absolute value of all correlations between block variables kept below r = 0.3.

After each block, participants answered two questions about how they perceived the block: (1) “How stressful was the last round?” and (2) “How exciting was the last round?”. Participants respond to each question by moving a slider to indicate a score between 0 and 100.

#### Timings

Each action involved a time cost: foraging took 1.5s, checking took 0.5s, hiding took 0.5s, and returning from hiding took 2s. Pressing one button inactivated all other buttons for the duration of that action’s time cost.

### Questionnaires

In the online session, participants were asked for demographic information (age, gender, education level, English fluency, and visual acuity). They also completed previously validated questionnaires to measure psychiatric symptoms and traits: the Apathy Motivation Index (AMI)^63^, the State-Trait Inventory for Cognitive and Somatic Anxiety (STICSA)^64^, the Snaith-Hamilton Pleasure Scale (SHAPS)^65^, and the Obsessive Compulsive Inventory (Revised) ‘Checking’ subscale (OCI-RC)^66^. One question from each of the three AMI, SHAPS, and OCI-R questionnaires was repeated as an indicator of consistency.

After the experimental session, participants completed a 29-item debrief questionnaire in which they reported their metacognitive awareness of their task behavior. They responded to a series of statements about the task by marking how often each statement was true for them. Response options were presented as a seven-point scale from 0 (‘Never’) to 6 (‘Very often’).

### Behavioral Analysis

We analysed participants’ behavior as a function of contextual factors. For button presses occurring before the predator had been discovered (‘pre-discovery phase’), we analysed the probability that each action was a check versus a forage given a combination of contextual factors: reward magnitude, an indication of the amount of reward available that was always visible on-screen; and *time pressure*, a measure of threat level computed as in Equation 1:

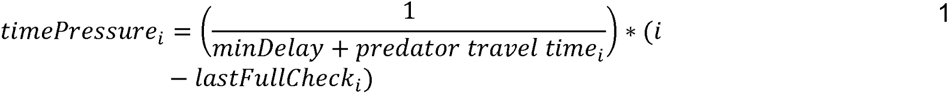

where *i* is the current time point, *minDelay* is the minimum possible delay for all predators in this task (2.5s), *predator travel time_i_* is the number of seconds that the predator type at *i* takes to reach the centre of the screen, and *lastFullCheck_i_* is the time point at which the participant most recently completed checking all areas of the environment (i.e., when they could be certain that no predator was present). For button presses occurring after the predator had been discovered (‘post-discovery phase’), we analysed 1) the probability that each action was a check or not, 2) the probability that each action was a forage or not, and 3) the probability that each action was a hide or not, based on three contextual factors: first, reward magnitude; second, *time since last check*, a measure of how long it had been since the participant had last seen the predator, computed as in Equation 2:

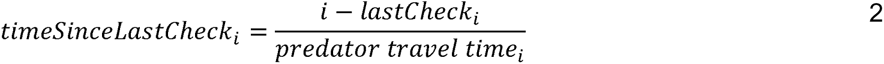

where *i* is the current time point, *lastCheck_i_* is the time point when the participant last saw the predator relative to the current time point *i*, and *predator travel time_i_* is again the time the current predator type takes to reach the centre of the screen at *i*; and third, *proximity*, a measure representing the amount of time until the predator would arrive, computed as in Equation 3:

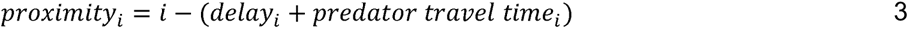

where *delay_i_* is the actual delay for the current predator and *i* and *predator travel time_i_* are as in Equations 1 and 2. Note that *proximity* is coded as a negative number, so that as the predator approaches, numbers become higher (i.e. less negative). Variables *time since last check* and *proximity* were both included because they provided measurements of different threat-related information: *proximity* was an expectation of predator arrival based on task knowledge that was not updated (a kind of heuristic), while *time since last check* was an estimate of threat level that was updated based on experience and thus more difficult to compute. All analyses of choice data were computed as non-hierarchical Bayesian regression models using the package brms{Citation} ^68,69^ with bernoulli(link=’logit’) link function. Regressions for both pre- and post-discovery phases were formulated as in Equation 4:

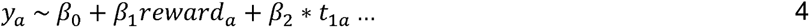

where *y_a_* is a binary variable indicating whether each action *a* in the relevant phase was the target action (either check, forage, or hide; post-PD actions analysed separately), *reward_a_* is the reward available at the time of action *a*, and *t_ia_* are the relevant threat-related variables computed for action *a* (see Table S1 for results and exact formulations). Regressors were z-score normalized and weak priors were set (normal distribution with mean 0 and standard deviation 3). Four chains were run with 4000 iterations and adapt_delta set to 0.8. Model fit was checked using Rhat < 1.1 and the absence of divergent samples. Model fits that did not meet these criteria were re-run with increased samples and adapt_delta. After computing each model non-hierarchically for each participant, we conducted two-tailed single-sample *t* tests across participants’ coefficient estimates to determine the parameter’s effect on choice data. Where outliers existed, test statistics are only reported as significant if the test was also significant with outliers excluded.

We also analysed participants’ inter-response times (IRTs) with respect to contextual factors such as action sequence. IRT was computed as the time in milliseconds between either the start of the block or the conclusion of the previous action (after the time cost associated with that action, when buttons were inactivated) and the next action. During analysis we found that participants often pressed buttons while they were inactivated (during the time cost of the most recent action) and we included these ‘inactive’ button presses in our IRT analyses. To the raw IRT values we applied a within-participant min-max transformation and across-participant outlier removal: Outlier values lower than the 25^th^ quartile minus 1.5 times the interquartile range (IQR) and greater than the 75^th^ quartile plus

1.5 times IQR across all participants were omitted. These analyses were computed as non-hierarchical Bayesian regressions using the package brms^67,68^ with ‘shifted_lognormal’ link function. Regressions were formulated as in Equation 5:

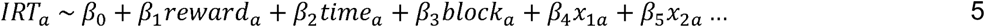

where reward_a_ is the current reward magnitude at action *a*, *time_a_* is the amount of time elapsed in the experiment at action *a*, *block_a_* is the block index for action *a*, and *x_is_* are contextual variables specific to each model (e.g., behavioral switch type) for action *a*. Modelling settings, model fit criteria, and parameter testing methods were the same as those used for the button press analysis (above).

### Neural Recording and Analysis

We used ultra-high field functional magnetic resonance imaging (7T fMRI) to identify brain activity corresponding to task behavior.

#### Data acquisition

We used a Siemens 7T MRI scanner to collect structural and functional MRI. High-resolution functional data were acquired using a multiband gradient-echo T2* echo planar imaging (EPI) sequence with a 1.5 x 1.5 x 1.5mm resolution; multiband acceleration factor 3; repetition time (TR) 1962ms; echo time (TE) 20ms; flip angle 68°; and a GRAPPA acceleration factor 2. Field of view (FOV) covered the whole brain with axial orientation and a fixed angulation of -30° (anterior-to-posterior phase encoding direction; 96 slices). In addition, a single-measurement, whole-brain, functional image with similar orientation (expanded functional image) was acquired prior to the main functional image and later used to improve registration of the main functional image. Structural data were acquired with a T1-weighted MP-RAGE sequence with a 0.7 x 0.7 x 0.7mm resolution; GRAPPA acceleration factor 2; TR 2200ms; TE 3.02ms; and inversion time (TI) 1050ms. To correct for magnetic field inhomogeneities a field map was acquired with a 2 x 2 x 2mm resolution; TR 620ms; TE1 4.08ms; TE2 5.10ms. Finally, cardiac and respiratory measurements were collected using pulse oximetry and respiratory bellows to regress out the effect of physiological noise in the functional data.

#### Data processing

Pre-processing was carried out using tools from FMRIB Software Library (FSL)^69–71^. Functional images were normalised, spatially smoothed (Gaussian kernel with 3mm full-width half-maximum), and temporally high-pass filtered (cut-off of 100s). Motion correction was performed using MCFLIRT^72^ and separation of brain from non-brain matter was performed using the Brain Extraction Tool (BET)^73^. Registration of functional images into Montreal Neurological Institute (MNI) space was carried out in three stages: first, the main functional image was registered to the expanded functional image using FMRIB’s Linear Image Registration Tool^72,74^ with three degrees of freedom (translation only); second, the main functional image was registered to the individual structural image using Boundary-Based Registration (BBR)^75^, incorporating Fieldmap correction; and third, the individual structural image was registered to standard space by using FMRIB’s Non-linear Image Registration Tool (FNIRT)^76^.

#### Whole-brain analyses

Whole-brain statistical analyses were performed at two-levels as implemented in FSL FEAT^77,78^. At the first (individual) level, we used a univariate general linear model (GLM) framework for each participant to estimate parameters. To account for temporal autocorrelations, first-level data were pre-whitened before group-level analysis^77^. The contrast of parameter and variance estimates from each participant were then combined at the second (group) level in a mixed-effects analysis (FLAME 1+2). The results were cluster-corrected with the voxel inclusion threshold *Z*=3.1 and cluster significance threshold of p<0.0002 two-tailed.

First-level analyses searched across the whole brain for voxels in which BOLD signal was associated with parametric variation in model variables. Our analysis split the time from the start of the delay period to the arrival of each predator (‘predator epochs’) into two phases: before the predator was discovered (‘pre-predator discovery’, or pre-PD phase) and after (‘post-predator discovery’ or post-PD phase). Model variables (see Figure S4 for correlation matrix) were computed separately for each phase. In the whole-brain analyses, a single GLM was used across the whole task. In the ROI analyses (see below), pre-PD and post-PD phases were analysed using separate GLMs. Importantly, in the whole-brain analysis, beyond the regressors detailed in the equations (GLM1, GLM2) below that were time-locked to action transitions, we also controlled for all other actions (all button presses related to foraging, checking and hiding, with forages and pre-PD checks segmented into first and subsequent actions), epoch threat type indexed by predator speed (slow, medium, and fast), and predator speed at the moment of predator discovery (see Figure S4).

For the pre-PD phase (GLM1), model variables included (see behavioral regressions): *reward* and *time pressure* (see Eq. 1), formulated as in Equation 6:

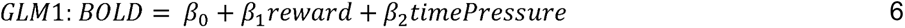

where *BOLD* is a column vector of time series data for a given voxel time-locked to a behavioral switch. For the post-discovery phase a very similar model (GLM2) was used but now, after seeing the predator, participants had access to an estimate of its proximity and so model variables included: *reward*; *time since last check* (see Eq. 2), and *proximity* (see Eq. 3), formulated as in Equation *7*:

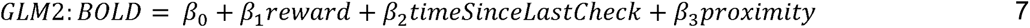

where *BOLD* is again voxel time series data time-locked to a behavioral switch. Variables *time since last check* and *proximity* were both included because they measured distinct threat-related information (correlated at *r* = 0.45 for switches to checking, *r* = 0.41 for switches to foraging, and *r* = 0.04 for switches to hiding; see correlations in Fig. S4). Regressors were modelled as stick functions (i.e., duration of zero), convoluted with a double-gamma hemodynamic response function (HRF). To reduce noise in the BOLD signal we added several task-unrelated confound regressors, including head motion parameters estimated by MCFLIRT in pre-processing; voxel-wise regressors created by physiological noise modelling (PNM)^79^ to model the effects of cardiac and respiratory noise; and regressors to remove timepoints corrupted by large motion that could not be corrected with MCFLIRT (across participants, 4±1% of timepoints were marked as corrupted by large motion).

Whole-brain analyses were time-locked to actions that represented behavioral switches (e.g., the first check after a series of forages). Time-locking to behavioural switches allowed enough time to separate events for analysis. Effect Required statistics provided by lower-level FSL FEAT^77^, a measure of each contrast’s efficiency/estimability, indicated that the average BOLD percent signal change required for any contrast of interest in the pre-discovery phase across all participants was 1.83 ± 0.58 (maximum 2.595). Due to participants conducting fewer checks after discovering a predator, contrasts in the post-discovery phase were more difficult to estimate; average BOLD percent signal change required across post-discovery contrasts was 2.46 ± 0.91 (maximum 3.64). Consequently, tests of post-discovery phase data (below) used only data from participants who checked >40 times and whose first-level statistics reported Effect Required below 2% for post-discovery check switch contrasts (N=13) or post-discovery forage switch contrasts (N=12).

### ROI time course analyses

To study the activity of regions of interest (ROIs), anatomical masks were created for each ROI in the MNI standard space using a conversion of the Talairach structural atlas (transformed into MNI space^80–82)^, mask templates from similar studies^23,83^, and cluster-corrected activations identified via whole-brain analysis. Next, masks were transformed from standard space to each participant’s structural space by applying a standard-to-structural warp field, transformed from structural to functional space by applying a structural-to-functional affine matrix, and binarised.

These masks were used to extract time-series data for analysis. First, a first-level whole-brain analysis was conducted for each participant with only regressors of no interest (all forages and all pre-discovery checks). Time-series data from each voxel within each ROI were then extracted from the residual functional data. Next, time-series data were averaged across the voxels within each ROI, normalised, up-sampled 20 times with cubic spline interpolation, and epoched in 17s windows starting from 2s before the button press to 15s after. Finally, GLMs were fit to each time step of the epoched data.

Given that the delay in hemodynamic response means that a BOLD signal change reflects neural activity ∼6s earlier^70^, we partitioned the ROI time courses into two phases for analysis: an early phase (0-5s post-action) when neural activity associated with the action of interest first becomes observable, and a late phase (5-10s post-action) that may reflect secondary neural processes associated with the action of interest. In support of this, a two-way repeated-measures ANOVA showed a main effect of phase on group peaks [*F*(1, 132) = 11.20, *p* = 0.001, ☐*_p_^2^* = 1.44] such that parametric variation in *time pressure* was linked with greater BOLD response in the early phase (*M* = 0.10, *SE* = 0.01) than in the later phase (*M* = 0.04, *SE* = 0.01) across VTA, SN, and DRN ROIs while controlling for time elapsed. Some tests were carried out on only the early or late phase depending on the expected timing of the effect: the early phase was used for analyses expected to reveal effects of action preparation and/or commission, while the late phase was used for analyses expected to reveal effects of action outcome.

For each psychophysiological interaction (PPI) analysis, time courses were time-locked to action switches in the pre-discovery phase and regressions were formulated as in Equation 8:

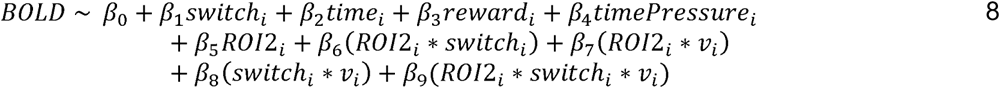

where *BOLD* is a *i* x *t* (*i* button press action, *t* time samples) matrix containing the time-series data from a given ROI during both switching to check and switching to forage; *switch_i_*is a binary variable indicating the behavioral switch (switch to check or forage) associated with action *i*, with the switch of interest coded positively; *time_i_* is the time elapsed in the experiment at action *i*; *reward_i_* is the reward magnitude available at the time of action *i*; *time pressure_i_*is a measure of threat level at action *i* (see Eq. 1); *ROI2* is the time course for another area of interest at action *i*; and *v_i_* is either *reward* or *time pressure* from action *i* depending on the effect of interest. To test for significance, we searched for peaks (or troughs) in each phase (early and late; see above) using a leave-one-out procedure to avoid any temporal selection biases: for a parameter of interest, a *β* weight value was selected for each participant from the time point identified as the peak average signal for the group minus that participant (similar to the approach used in Khalighinejad et al.^23^). Selected values were tested via two-tailed single sample *t* tests. Further correction for multiple comparisons was considered unnecessary because ROIs were chosen based on their significance in cluster-corrected whole-brain analysis (p<0.0002 two-tailed), which itself performs rigorous correction for multiple comparisons.

To test for overall effects of factors such as ROI location (cortical versus sub-cortical) and behavioral switch type on functional connectivity, we performed ANOVAs on fitted parameter peak values selected using the same method. Peaks from both the early time window (-2-5s, capturing effects theoretically associated with action preparation) and late time window (5-10s, capture effects associated with action commission) were included in ANOVAs.

#### Replication analyses

All the previous analyses focused on activity in distributed neural circuits linked to decisions to forage for rewards or check for threats in the pre-PD task phase. We initially focused on this task phase because it contained similar levels of checks and forages. In the final stage of the analysis, however, we examined whether we could replicate findings from the pre-PD phase in the post-PD phase. It was not possible to analyze the data for all participants in the post-PD phase because some only made a very small number of checks during this task phase. When testing the possibility that we could replicate results (shown in Fig. 7A, C, D), we therefore focused on participants who had made more checks (>40) in the post-PD phase and in whom an analysis of effects sizes indicated <2% change in BOLD signal was required to detect an effect of the parameter of interest (threat level or reward) during switches to check (leaving N=12 in all replication analyses). We took an analogous approach when testing the replicability of the results in Figure 6 albeit now focusing on participants in whom an analysis of effects sizes indicated <2% change in BOLD signal was required to detect an effect of the parameter of interest during switches to forage (leaving N=12 for replication analyses related to foraging and N=13 for those related to checking).

To test whether key findings from our fMRI analysis replicated, we regressed variables of interest against ROI time course data with models formulated as in Equation 9:

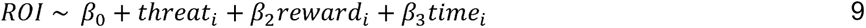

where *ROI* is the time course data processed as in the PPI analyses; *threat_i_* is a measure of threat level at action *I* as indexed by either *time pressure* or *proximity* for the pre- and post-discovery phases respectively; *reward_i_* is the reward available at action *I*; and *time_i_* is the time elapsed at action *i*. Models were fit for each ROI in the distributed neural circuits presented in the main text (ACC, AI, DRN, LHb, SN, and VTA), with separate models for the pre- and post-discovery phase data. Peaks were selected from parameters fit to pre-discovery phase data using the leave-one-out procedure (above). Here the peak search was constrained to a time window dictated by significant PPI results reported in Table S17: For tests involving ROIs in the threat-related circuit, we used the range of mean peak times reported for all PPIs associated with action commission within that circuit; Instead of searching for PPI effects occurring at the exact same time as the pre-PD phase, we used this peak search procedure to slightly expand the search window to allow for task-related differences between the pre- and post-PD phases: due to fewer repeated checks in the post-PD phase, the timing of behavioral switch effects might be expected to be slightly different. Selected peaks were tested with two-tailed single sample t-tests. Tests significant at the *p* < 0.05 level were then replicated using the equivalent parameters fit to post-discovery phase data. Peaks were selected from these parameters by searching within the time range that contained 95% of the selected peaks from the equivalent pre-discovery phase test. Selected peaks were then tested with a one-tailed single-sample t-test, with the tail corresponding to the direction of effect in the pre-discovery phase test. Results of replication tests are in Table S9.

## Supporting information

Task demo

## Acknowledgements

Supported by a Wellcome Trust grant (221794/Z/20/Z), BBSRC Discovery Fellowships (BB/W008947/1 and BB/V004999/1), an MRC Skills Development Fellowship (MR/N014448/1), and the Institut National de la Santé et de la Recherche Médicale.

