## Supplemental Materials for "An ancient subcortical circuit decides when to orient to threat in humans"

### Supplementary Figures

Figure S1. **Related to Figure 4Ai.** Interactions between activity in cortical areas ACC and AI and subcortical areas Hb and DRN as a function of transitioning to checking (A-C) and as a function of both transitioning to checking and threat level (as indexed by time pressure) (D -F). Peak values in A-C were entered into the two-way ANOVA summarized in Fig. 4Ai and reported in Table S6. Peak values in D-F were entered into the two-way ANOVA summarized in Fig. 4Bi and reported in Table S6. In general, all the interaction patterns shown here, in contrast to those shown in Fig. 4Aii and Fig. 4Bii are non-significant.

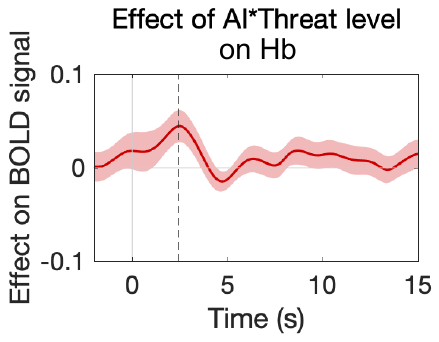

Figure S2. Early phase Hb activity was moderated by a significant interaction between threat level (as indexed by time pressure) and AI activity (p < 0.05).

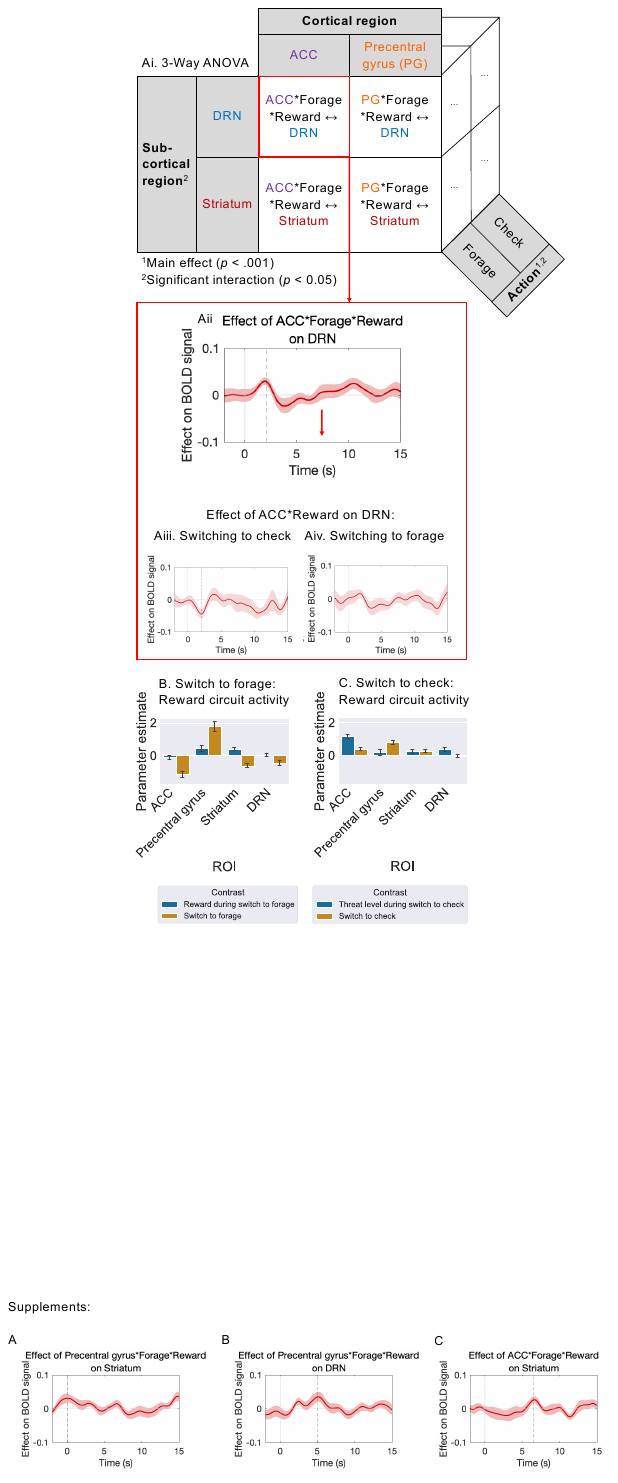

Figure S3. **Related to Figure 6** Interactions between activity in precentral gyrus and striatum as a function of transitioning to foraging and reward level. A three-way interaction between precentral gyrus activity, switching to forage, and reward level moderated activity in the striatum (both p < 0.05). PPI statistics reported in Table S17.

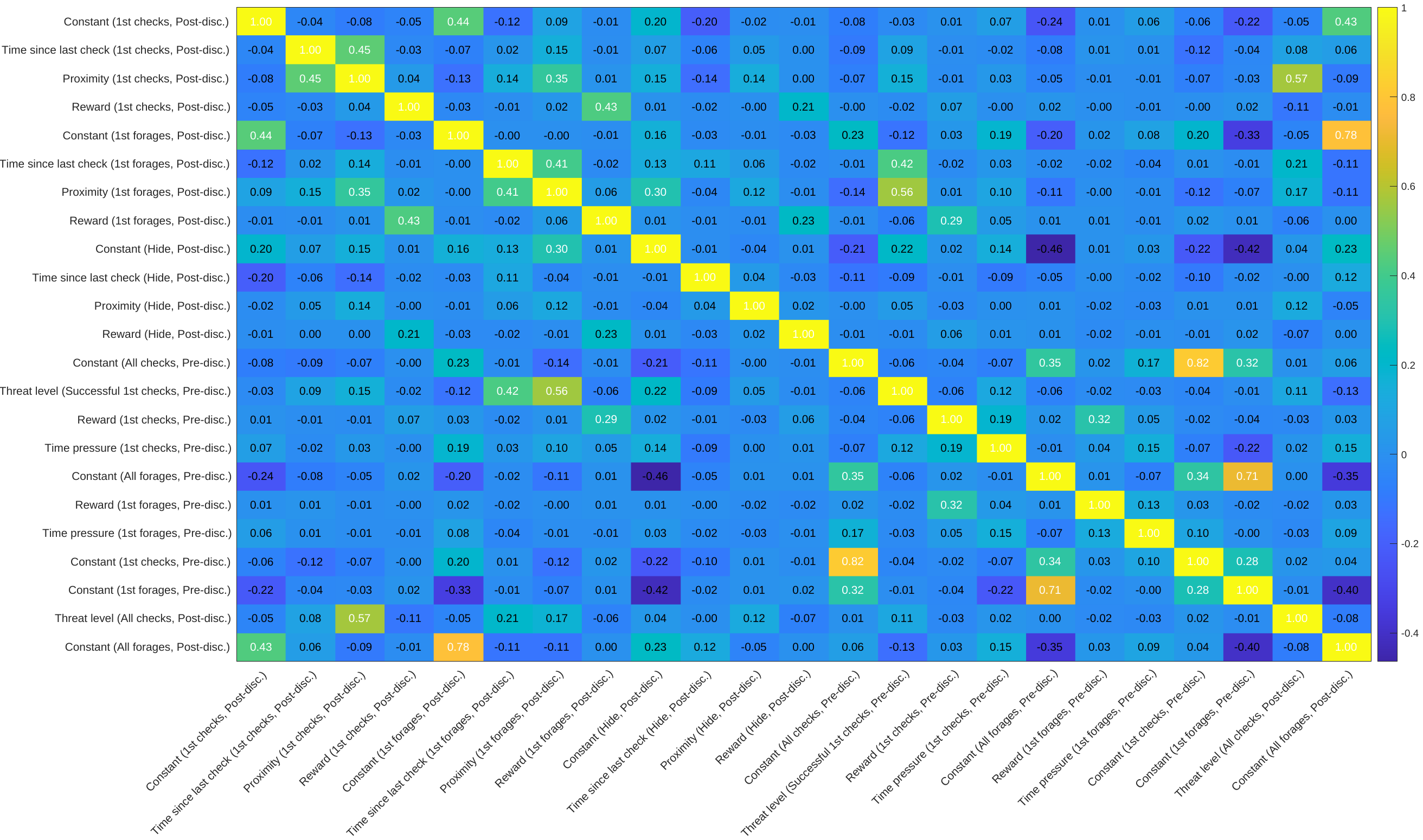

*Figure S4.* **Correlation matrix for regressors included in the fMRI analysis, averaged across all participants.** Constant terms from the regression model denoting the first check in a series of checks and the subsequent checks in the same series were correlated as were the first forage and subsequent forages in a series of forages. However, there were no other correlations affecting the interpretation of key analyses.

### Supplementary Tables

#### Table S1.

*Two-Tailed Single Sample T-Tests Across Regression Coefficients Showing Effects of Contextual Factors on Action Selection (Related to Fig. 1H)*

| Parameter | *P* value | T statistic | 95% CI | | Degrees of freedom | Mean | SD |
| --- | --- | --- | --- | --- | --- | --- | --- |
|  |  |  | Lower | Upper |  |  |  |
| Reward^1^ | 0 | -5.38 | -0.39 | -0.17 | 22 | -0.28 | 0.25 |
| Time pressure^1^ | 0 | 12.94 | 1.18 | 1.64 | 22 | 1.41 | 0.52 |
| Reward^2^ | 0.242 | -1.2 | -0.19 | 0.05 | 22 | -0.07 | 0.27 |
| Time since last check^2^ | 0 | 5.24 | 3.24 | 7.49 | 22 | 5.37 | 4.91 |
| Proximity^2^ | 0 | -7.72 | -0.15 | -0.09 | 22 | -0.12 | 0.08 |
| Reward^3^ | 0.256 | 1.17 | -0.03 | 0.09 | 22 | 0.03 | 0.14 |
| Time since last check^3^ | 0 | -8.72 | -7.02 | -4.32 | 22 | -5.67 | 3.12 |
| Proximity^3^ | 0 | -14.61 | -0.43 | -0.32 | 22 | -0.37 | 0.12 |
| Reward^4^ | 0.972 | 0.04 | -0.07 | 0.07 | 22 | 0 | 0.16 |
| Time since last check^4^ | 0.018 | 2.56 | 0.37 | 3.56 | 22 | 1.97 | 3.68 |
| Proximity^4^ | 0 | 16.77 | 0.67 | 0.86 | 22 | 0.76 | 0.22 |

^1^Model: $Check (Pre disc. phase) \sim1+ \beta_{1}*Reward+ \beta_{2}*TimePressure$

^2^Model: $Check \left( Post disc. phase \right)\sim1+ \beta_{1}*Reward+ \beta_{2}*TimeSinceLastCheck+ \beta_{3}*Proximity$

^3^Model: $Forage \left( Post disc. phase \right)\sim1+ \beta_{1}*Reward+ \beta_{2}*TimeSinceLastCheck+ \beta_{3}*Proximity$

^4^Model: $Hide \left( Post disc. phase \right)\sim1+ \beta_{1}*Reward+ \beta_{2}*TimeSinceLastCheck+ \beta_{3}*Proximity$

#### Table S2.

*Two-Tailed Single Sample T-Tests Across Regression Coefficients Showing Effects of Transition, Action Type, and Contextual Factors on Inter-Response Times (IRTs) (Related to Fig. 1I)*

| Parameter | *P* value | T statistic | 95% CI | | Degrees of freedom | Mean | SD |
| --- | --- | --- | --- | --- | --- | --- | --- |
|  |  |  | Lower | Upper |  |  |  |
| Transition | 0 | 10.16 | 0.18 | 0.27 | 22 | 0.22 | 0.1 |
| Check | 0 | 4.23 | 0.14 | 0.41 | 22 | 0.27 | 0.31 |
| Transition*Check | 0 | -8.07 | -0.31 | -0.18 | 22 | -0.24 | 0.14 |
| *Time pressure* | 0 | 7.3 | 0.13 | 0.23 | 22 | 0.18 | 0.12 |
| *Reward* | 0 | -4.56 | -0.06 | -0.02 | 22 | -0.04 | 0.04 |
| Time elapsed | 0 | -6.94 | -0.06 | -0.03 | 22 | -0.05 | 0.03 |
| Block | 0.043 | -2.15 | -0.02 | 0 | 22 | -0.01 | 0.02 |

*Note.* Model: $RT \left( Pre disc. phase \right)\sim1+ \beta_{1}*transition+ \beta_{2}*check+\beta_{3}*(transition*check)+\beta_{4}*timePressure+\beta_{5}*reward+\beta_{6}*timeElapsed+\beta_{7}*block$

#### Table S3.

*Two-Tailed Single Sample T-Tests Across Regression Coefficients Showing Effects of Approaching a Switch to Checking on Inter-Response Times (IRTs) between Forages (Related to Fig. 1K)*

| Parameter | *P* value | T statistic | 95% CI | | Degrees of freedom | Mean | SD |
| --- | --- | --- | --- | --- | --- | --- | --- |
|  |  |  | Lower | Upper |  |  |  |
| *Time pressure* | 0 | 7.24 | 0.2 | 0.37 | 22 | 0.28 | 0.19 |
| *Reward* | 0 | -5.94 | -0.12 | -0.06 | 22 | -0.09 | 0.07 |
| Time elapsed | 0 | -5.22 | -0.1 | -0.04 | 22 | -0.07 | 0.06 |
| Proximity to check switch | 0 | 4.4 | 0.06 | 0.15 | 22 | 0.1 | 0.11 |
| Block | 0.156 | -1.47 | -0.02 | 0 | 22 | -0.01 | 0.03 |

*Note.* Model: $RT \left( Pre disc. phase \right)\sim1+ \beta_{1}\mathrm{timePressure}+ \beta_{2}reward+\beta_{3}timeElapsed+\beta_{4}proxToCheckSwitch+\beta_{5}*block$

#### Table S4.

*Two-Tailed Single Sample T-Tests Across Regression Coefficients Showing Effects of Predator Discovery on Inter-Response Times (IRTs) (Related to Fig. 1L)*

| Parameter | *P* value | T statistic | 95% CI | | Degrees of freedom | Mean | SD |
| --- | --- | --- | --- | --- | --- | --- | --- |
|  |  |  | Lower | Upper |  |  |  |
| First action post-predator discovery | 0 | 6.18 | 0.07 | 0.13 | 22 | 0.1 | 0.08 |
| *Reward* | 0.011 | -2.79 | -0.06 | -0.01 | 22 | -0.04 | 0.06 |
| Block | 0.03 | -2.32 | -0.02 | 0 | 22 | -0.01 | 0.02 |
| Time elapsed | 0 | -5.92 | -0.05 | -0.03 | 22 | -0.04 | 0.03 |
| Pre-discovery phase (vs. post) | 0.004 | -3.17 | -0.08 | -0.02 | 22 | -0.05 | 0.08 |

*Note.* Model: $RT \left( Pre+post disc. phases \right)\sim1+ \beta_{1}*postPredatorDisc+ \beta_{2}*reward+\beta_{3}*block+\beta_{4}*timeElapsed+\beta_{5}*preDiscPhase$

#### Table S5.

*Whole-Brain Cluster-Corrected Activations and their Peak Values within ROIs*

| Time-locked action | Parameter | Cluster type | ROI | Peak MNI coord. | Peak Z | No. of voxels* | Cluster index |
| --- | --- | --- | --- | --- | --- | --- | --- |
| Check switch | Constant | Act. | Pulvinar | (-14, -22, 12) | 4.44 | 38 | 1 |
| Check switch | Constant | Act. | Striatum | (-16, 14, -2) | 3.67 | 4 | 3 |
| Check switch | Constant | Act. | ACC | (-2, 12, 32) | 3.94 | 21 | 7 |
| Check switch | Constant | Act. | Precentral gyrus | (-40, -24, 64) | 4.54 | 23 | 7 |
| Check switch | Threat level | Act. | ACC | (0, 16, 28) | 5.30 | 95 | 2 |
| Check switch | Threat level | Act. | DRN | (2, -30, -16) | 4.43 | 22 | 2 |
| Check switch | Threat level | Act. | Hb | (-4, -24, 0) | 4.35 | 26 | 2 |
| Check switch | Threat level | Act. | AI | (30, 22, -10) | 6.00 | 422 | 2 |
| Check switch | Threat level | Act. | PAG | (6, -30, -10) | 4.28 | 37 | 2 |
| Check switch | Threat level | Act. | Pulvinar | (8, -24, 14) | 4.81 | 88 | 2 |
| Check switch | Threat level | Act. | SC | (4, -28, -4) | 4.61 | 50 | 2 |
| Check switch | Threat level | Act. | SN | (-6, -20, -12) | 4.19 | 57 | 2 |
| Check switch | Threat level | Act. | Striatum | (-14, 18, 4) | 4.18 | 24 | 2 |
| Check switch | Threat level | Act. | VTA | (-4, -20, -14) | 3.86 | 5 | 2 |
| Forage switch | Constant | Act. | Precentral gyrus | (-40, -30, 64) | 6.19 | 44 | 4 |
| Forage switch | Constant | Deact. | Striatum | (8, 12, 2) | 6.17 | 168 | 9 |
| Forage switch | Constant | Deact. | DRN | (0, -32, -20) | 4.46 | 34 | 12 |
| Forage switch | Constant | Deact. | Hb | (-4, -24, 0) | 4.97 | 31 | 12 |
| Forage switch | Constant | Deact. | PAG | (4, -30, -6) | 4.65 | 48 | 12 |
| Forage switch | Constant | Deact. | Pulvinar | (10, -24, 0) | 4.69 | 15 | 12 |
| Forage switch | Constant | Deact. | SC | (-4, -30, -4) | 5.51 | 77 | 12 |
| Forage switch | Constant | Deact. | SN | (8, -26, -16) | 4.26 | 58 | 12 |
| Forage switch | Constant | Deact. | VTA | (-4, -12, -10) | 3.76 | 6 | 12 |
| Forage switch | Constant | Deact. | AI | (-30, 20, 10) | 6.11 | 419 | 13 |
| Forage switch | Constant | Deact. | ACC | (-2, 16, 24) | 5.41 | 97 | 14 |
| Forage switch | Threat level | Act. | Hb | (6, -24, 0) | 3.74 | 6 | 1 |
| Forage switch | Threat level | Act. | PAG | (4, -30, -10) | 4.24 | 18 | 1 |
| Forage switch | Threat level | Act. | Pulvinar | (10, -24, 0) | 3.92 | 3 | 1 |
| Forage switch | Threat level | Act. | SC | (4, -30, -4) | 3.92 | 13 | 1 |
| Forage switch | Threat level | Act. | AI | (32, 28, 4) | 4.78 | 144 | 8 |
| Forage switch | Threat level | Deact. | Striatum | (10, 12, -4) | 4.54 | 57 | 6 |
| Forage switch | Reward | Act. | Striatum | (-12, 14, 4) | 4.24 | 63 | 2 |
| Forage switch | Reward | Act. | Precentral gyrus | (-40, -24, 64) | 3.85 | 8 | 3 |
| Forage switch | Pre-disc. reward plus post-disc. reward | Act. | Striatum | (-16, 10, 0) | 4.51 | 199 | 3 |
| Forage switch | Pre-disc. reward plus post-disc. reward | Act. | Precentral gyrus | (-40, -22, 64) | 3.93 | 10 | 5 |

*Note.* All Z > 3.1, *P* < 0.0001. All time-locked actions are from the pre-discovery phase unless otherwise specified.

*Where the cluster activation/deactivation overlaps with the ROI.

#### Table S6.

*ROI Statistics from Whole-Brain Analysis Extracted for Each Participant*

See attached TableS6.csv.

#### Table S7.

*PPI Analyses: Two-Tailed Single Sample T-Tests Across Regression Coefficients*

See attached TableS7.csv.

*Average time of peaks across which single sample *t* test was significant, relative to button press onset.

^†^Model formulas:

Model 1: $ROI time course \left( check switch and forage switch, pre disc. phase \right) \sim\beta_{0}+ \beta_{1}Check+\beta_{2}TimePressure+\beta_{3}*eward+\beta_{4}Time+\beta_{5}+ROI2+\beta_{6}\left( ROI2*Check \right)+\beta_{7}\left( ROI2*TimePressure \right)+\beta_{8}\left( Check*TimePressure \right)+\beta_{9}\left( ROI2*Check*TimePressure \right)$

Model 2: $ROI time course \left( Successful check switch and forage switch, pre disc. phase \right) \sim\beta_{0}+ \beta_{1}Check+\beta_{2}TimePressure+{\beta_{3}Reward+\beta}_{4}Time+\beta_{5}+ROI2+\beta_{6}\left( ROI2*Check \right)+\beta_{7}\left( ROI2*TimePressure \right)+\beta_{8}\left( Check*TimePressure \right)+\beta_{9}(ROI2*Check*TimePressure)$

Model 3:

$$ROI time course \left( Unsuccessful check switch and forage switch, pre disc. phase \right) \sim\beta_{0}+ \beta_{1}Check+\beta_{2}TimePressure+\beta_{3}Reward+\beta_{4}Time+\beta_{5}+ROI2+\beta_{6}\left( ROI2*Check \right)+\beta_{7}\left( ROI2*TimePressure \right)+\beta_{8}\left( Check*TimePressure \right)+\beta_{9}(ROI2*Check*TimePressure)$$

Model 4:

$$ROI time course \left( check switch and forage switch, pre disc. phase \right)\sim\beta_{0}+ \beta_{1}Forage+\beta_{2}Reward+\beta_{3}TimePressure+\beta_{4}Time+\beta_{5}+ROI2+\beta_{6}\left( ROI2*Forage \right)+\beta_{7}\left( ROI2*Reward \right)+\beta_{8}\left( Forage*Reward \right)+\beta_{9}(ROI2*Forage*Reward)$$

#### Table S8.

*Post-Hoc Tukey HSD Test To Examine the Main Effect of Cortical ROI in ANOVA Reported in* Table S6 *(Related to Fig. 4Ai)*

| Group1 | Group2 | Meandiff | p-adj | Lower | Upper | Reject |
| --- | --- | --- | --- | --- | --- | --- |
| ACC | AI | -0.0232 | 0.0117 | -0.0411 | -0.0052 | True |

#### Table S9.

*Two-Way ANOVA Testing the Effects of Cortical ROI and Sub-cortical ROI on the Extent to which Functional Connectivity is Moderated by Switching to Check (Related to Fig. 4Ai)*

|  | sum_sq | df | F | PR(>F) |
| --- | --- | --- | --- | --- |
| C(Cortical ROI) | 0.024722 | 1.0 | 7.032443 | 0.008719 |
| C(Sub-cortical ROI) | 0.004118 | 1.0 | 1.171459 | 0.280549 |
| C(Cortical ROI):C(Sub-cortical ROI) | 0.056697 | 1.0 | 16.128202 | 0.000087 |
| Residual | 0.632770 | 180.0 | NaN | NaN |

*Note.* Cortical areas included ACC and AI and subcortical areas included Hb and DRN. Data entered into the ANOVA were both early and late peaks selected (using a leave-one-out procedure on the group signal) from parameters fit to $\beta_{6}$ from the following regression:

$$Subcortical ROI time course (Check switch and forage switch, pre disc. phase\sim\beta_{0}+ \beta_{1}SwitchToCheck+\beta_{2}TimePressure+\beta_{3}Time+\beta_{4}Reward+\beta_{5}+CorticalROI+\beta_{6}\left( CorticalROI*SwitchToCheck \right)+\beta_{7}\left( CorticalROI*TimePressure \right)+\beta_{8}\left( SwitchToCheck*TimePressure \right)+\beta_{9}(CorticalROI*SwitchToCheck*TimePressure)$$

#### Table S10.

*Two-Way ANOVA Testing the Effects of Cortical ROI and Sub-cortical ROI on the Extent to which Functional Connectivity is Moderated by Switching to Check and Threat Level (Related to Fig. 4Bi)*

|  | sum_sq | df | F | PR(>F) |
| --- | --- | --- | --- | --- |
| C(Cortical ROI) | 0.023501 | 1.0 | 4.229004 | 0.041182 |
| C(Sub-cortical ROI) | 0.061755 | 1.0 | 11.113015 | 0.001041 |
| C(Cortical ROI):C(Sub-cortical ROI) | 0.004722 | 1.0 | 0.849757 | 0.357855 |
| Residual | 1.000259 | 180.0 | NaN | NaN |

*Note.* Cortical areas included ACC and AI and subcortical areas included Hb and DRN. Data entered into the ANOVA were both early and late peaks selected (using a leave-one-out procedure on the group signal) from parameters fit to $\beta_{9}$ from the following regression:

$$Subcortical ROI time course \left( First checks and first forages, pre disc. phase \right)\sim\beta_{0}+ \beta_{1}*SwitchToCheck+\beta_{2}*TimePressure+\beta_{3}*Time+\beta_{4}*Reward+\beta_{5}+CorticalROI+\beta_{6}*\left( CorticalROI*SwitchToCheck \right)+\beta_{7}*\left( CorticalROI*TimePressure \right)+\beta_{8}*\left( SwitchToCheck*TimePressure \right)+\beta_{9}*(CorticalROI*SwitchToCheck*TimePressure)$$

#### Table S11.

*Post-Hoc Tukey HSD Test to Examine the Main Effect of Sub-Cortical ROI in ANOVA Reported in* Table S8 *(Related to Fig. 4Bi)*

| Group1 | Group2 | Meandiff | p-adj | Lower | Upper | Reject |
| --- | --- | --- | --- | --- | --- | --- |
| DRN | HB | -0.0366 | 0.0011 | -0.0585 | -0.0148 | True |

#### Table S12.

*Post-Hoc Tukey HSD Test to Examine the Main Effect of Cortical ROI in ANOVA Reported in* Table S8 *(Related to Fig. 4Bi)*

| Group1 | Group2 | Meandiff | p-adj | Lower | Upper | Reject |
| --- | --- | --- | --- | --- | --- | --- |
| ACC | AI | 0.0226 | 0.0467 | 0.0003 | 0.0449 | True |

#### Table S13.

*Two-Way ANOVA Testing the Effects of Action Type (Switch to Forage vs. Switch to Check) and ROI on the Extent to which Hb Activity is Moderated By ROI, Action, and Time Pressure (Related to Fig. 5B-D)*

|  | sum_sq | df | F | PR(>F) |
| --- | --- | --- | --- | --- |
| **C(ROI)** | 0.000 | 2.000 | 0.000 | 1.000 |
| **C(Action)** | 0.020 | 1.000 | 4.942 | 0.027 |
| **C(ROI):C(Action)** | 0.022 | 2.000 | 2.741 | 0.066 |
| **Residual** | 1.102 | 270.000 | NaN | NaN |

*Note.* ROIs included SN, VTA, and DRN and actions included switching to check and switching to forage. Data entered into the ANOVA were both early and late peaks selected (using a leave-one-out procedure on the group signal) from parameters fit to $\beta_{9}$ from the following regression:

$$Hb time course \left( First checks and first forages, pre disc. phase \right)\sim\beta_{0}+ \beta_{1}*Action+\beta_{2}*TimePressure+\beta_{3}*Time+\beta_{4}*Reward+\beta_{5}+ROI+\beta_{6}*\left( ROI*Action \right)+\beta_{7}*\left( ROI*TimePressure \right)+\beta_{8}*\left( Action*TimePressure \right)+\beta_{9}*(ROI*Action*TimePressure)$$

where *Hb* time course was BOLD signal time-locked to either a switch to forage or switch to check action (both models computed separately).

#### Table S14.

*Post-Hoc Tukey HSD Test to Examine the Main Effect of Action in ANOVA Reported in* Table S13 *(Related to Fig. 5Ai)*

| Group1 | Group2 | Meandiff | p-adj | Lower | Upper | Reject |
| --- | --- | --- | --- | --- | --- | --- |
| Switch to check | Switch to forage | 0.0358 | 0.0001 | 0.0177 | 0.0539 | True |

#### Table S15.

*Two-Way ANOVA Testing the Effects of Check Outcome and ROI on the Extent to which Hb Activity is Moderated by ROI, Switching to Check, and Threat Level (Related to Fig. 5E-J)*

|  | sum_sq | df | F | PR(>F) |
| --- | --- | --- | --- | --- |
| C(ROI) | 0.006630 | 2.0 | 0.194274 | 0.823667 |
| C(Outcome) | 0.405684 | 1.0 | 23.774707 | 0.000003 |
| C(ROI):C(Outcome) | 0.007568 | 2.0 | 0.221746 | 0.801416 |
| Residual | 2.252408 | 132.0 | NaN | NaN |

*Note.* ROIs included SN, VTA, and DRN and outcomes included discovering a new predator versus not seeing any predator. Data entered into the ANOVA were both early and late peaks selected (using a leave-one-out procedure on the group signal) from parameters fit to $\beta_{9}$ from the following regression:

$$Hb time course \left( First checks and first forages, pre disc. phase \right)\sim\beta_{0}+ \beta_{1}SwitchToCheck+\beta_{2}TimePressure+\beta_{3}Time+\beta_{4}Reward+\beta_{5}+ROI+\beta_{6}\left( ROI*SwitchToCheck \right)+\beta_{7}\left( ROI*TimePressure \right)+\beta_{8}\left( SwitchToCheck*TimePressure \right)+\beta_{9}(ROI*SwitchToCheck*TimePressure)$$

where *Hb* time course was BOLD signal time-locked to either a switch to forage or switch to check action (both models computed separately).

#### Table S16.

*Results from Replication Analyses Comparing the Effects of Key Variables on ROI Time Courses in Pre- Versus Post-Discovery Phases*

See attached TableS16.csv.

#### Table S17

*PPI Results: Two-Sided Wilcoxon Signed Rank Tests*

| ROI | Seed ROI | Regressor^2^ | *P* value | Z | Mean | SD | Mean peak time^1^ (s) | Figure |
| --- | --- | --- | --- | --- | --- | --- | --- | --- |
| Hb | ACC | ACC*Check | 0.001 | 3.28 | 0.05 | 0.06 | -0.88 | 4Aii |
| DRN | AI | AI*Threat*Check | 0.048 | 1.98 | 0.02 | 0.07 | 4.49 | 4Bii |
| DRN | AI | AI*Threat*Check | 0.024 | 2.25 | 0.04 | 0.07 | 4.98 | 4Bii |
| SN | Hb | Hb*Threat*Check (Successful outcome) | 0.015 | -2.43 | -0.10 | 0.20 | 8.61 | 5E |
| VTA | Hb | Hb*Threat*Check (Successful outcome) | 0.003 | -2.95 | -0.10 | 0.13 | 8.71 | 5F |
| DRN | Hb | Hb*Threat*Check (Successful outcome) | 0.036 | -2.10 | -0.08 | 0.19 | 7.43 | 5G |
| Hb | AI | AI*Threat | 0.031 | 2.16 | 0.04 | 0.08 | 2.42 | S2 |
| SN | Hb | Hb*Threat*Check | 0.031 | 2.16 | 0.03 | 0.06 | 6.35 | 5A |
| VTA | Hb | Hb*Threat*Check | 0.033 | 2.13 | 0.02 | 0.06 | 5.76 | 5B |
| DRN | Hb | Hb*Threat*Check | 0.042 | 2.04 | 0.02 | 0.05 | 5.27 | 5C |
| Striatum | Precentral gyrus | Precentral gyrus*Reward*Forage | 0.048 | 1.98 | 0.03 | 0.06 | 0.06 | S3 |

^1^Average time of peaks across which two-sided Wilcoxon signed rank test was significant, relative to button press onset.
